# Chromatin tethering to the nuclear envelope enhances its accessibility to RNAPII and promotes chromatin asymmetric organization

**DOI:** 10.64898/2026.03.06.710131

**Authors:** Dana Lorber, Ido Azuri, Amit Kumar, Ron Rotkopf, Sam Safran, Talila Volk

## Abstract

The intrinsic tendency of chromatin to self-attract competes with its association with RNA Polymerase II (RNAPII), a prerequisite for efficient transcription. Using high-resolution live imaging of chromatin and RNAPII organization in *Drosophila* larval muscle nuclei, we demonstrate that chromatin tethering to the nuclear envelope via the Linker of Nucleoskeleton and Cytoskeleton (LINC) complex is essential for maintaining chromatin three-dimensional organization. Disruption of chromatin-lamina interactions either in LINC mutants or following knockdown of Barrier-to-Autointegration Factor (BAF) results in enhanced chromatin clustering in the nucleoplasm and reduced RNAPII-chromatin interaction. Consistent with these observations, computer simulations revealed an inverse relationship between the chromatin cluster size and the degree of chromatin tethering to the nuclear lamina. We also measured chromatin distribution with respect to the nuclear lamina and found asymmetry of RNAPII distributions within chromatin clusters, which correlated with their proximity to the nuclear envelope, a relationship that is lost in nuclei lacking a functional LINC complex.

Our findings demonstrate that chromatin association with the nuclear envelope counteracts chromatin self-attraction and facilitates RNAPII binding to DNA.

## Introduction

Packaging of chromatin within the nucleus organizes the genome into three-dimensional structures that influence transcription regulation; at the same time, transcription itself reshapes chromatin architecture, creating a dynamic interplay between both processes (Misteli 2020; Dekker et al. 2024; Almassalha et al. 2025; Liu et al. 2023; Boettiger et al. 2016). To fit within the nucleus, chromatin is packaged in a hierarchical manner, from loops and compartments to chromosome territories (T. Cremer and Cremer 2001; Sanders et al. 2022; Elimelech and Birnbaum 2020). Within compartments, chromatin further organizes into clusters and clutches (Ricci et al. 2015; Szabo et al. 2018). These clusters consist of a dense chromatin core, corresponding to heterochromatin; the chromatin density decreases towards cluster periphery, which is dominated by less densely packed euchromatin (Kant et al. 2024; Y. Li et al. 2022; Marion Cremer et al. 2017; Gelléri, Márton et al. 2024; Miron et al. 2020; Nozaki et al. 2017), with the overall concentration ranging from 5 Mbp/µm^3^ up to 300 Mbp/µm^3^ (Gelléri et al. 2023). Chromatin near the periphery of the clusters undergoes continuous molecular exchange with soluble DNA-binding transcription complexes, including RNAPII (Goronzy et al. 2022; Ricci et al. 2015; W. Cho et al. 2016) and Transcription factor II H (Verschure et al. 2003). In addition, the chromatin interacts with nuclear organelles, e.g., nucleolus via sequences called Nucleolus-Associated Domains (NADs) (Peng et al. 2023; Bersaglieri and Santoro 2023) or the nuclear lamina via Lamina-Associated Domains (LADs) (van Steensel and Belmont 2017; Ulianov et al. 2019). LADs sequences represent about 30%-40% of the genome (Manzo, Dauban, and van Steensel 2022; van Bemmel et al. 2010), and are generally associated with repressive histone modifications such as H3K9me2/3 or H3K27me3 in a cell-type-specific manner (Guelen et al. 2008; Towbin et al. 2012; Padeken, Methot, and Gasser 2022).

Using live-cell imaging of *Drosophila* muscle cells, we previously demonstrated that chromatin is predominantly localized at the nuclear periphery, adjacent to the nuclear lamina. This arrangement results in a relatively dense chromatin layer near the inner nuclear membrane, leaving a substantial central nuclear volume largely devoid of chromatin (Amiad-Pavlov, Lorber et al. 2021). This spatial organization was suggested to arise from the balance between chromatin self-attraction and chromatin affinity for the nuclear lamina (Bajpai et al. 2021), and has been observed in other cell types as well (Amiad-Pavlov, Lorber et al. 2021). Importantly, live chromatin imaging was crucial for revealing its 3D peripheral organization (Lorber and Volk 2022), since fixation may induce chromatin aggregation (Dubochet and Blanc 2001), change the native state of liquid–liquid phase separation (Irgen-Gioro et al. 2022), increase chromatin compaction (Dupont et al. 2023), and reduce nucleosome mobility (Nozaki et al. 2023). The finer-scale organization of chromatin at the nuclear lamina and the functional significance of its peripheral organization remain poorly understood. Chromatin binding to the nuclear lamina is mediated by proteins associated with the inner nuclear membrane. One key example is the Barrier to Autointegration Factor (BAF), which binds directly to DNA (Sears and Roux 2020; Andrés and González 2009), as well as to LAP2, Emerin and MAN1 (LEM)-domain proteins at the nuclear membrane (Barton, Soshnev, and Geyer 2015; Gruenbaum and Foisner 2015; Sobo et al. 2024; Marcelot et al. 2021), reinforcing the connection between the genome and the nuclear periphery. BAF plays a pivotal role in nuclear function (Marcelot et al. 2021); it contributes to chromatin compaction, membrane reassembly following mitosis (Sears and Roux 2020), nuclear envelope repair during interphase (Sears and Roux 2020; Marcelot et al. 2021), and DNA break sites (Sears and Roux 2020).

Another major complex that mediates chromatin interaction with the nuclear envelope is the Linker of Nucleoskeleton and Cytoskeleton (LINC) complex that couples cytoskeletal elements and the nucleoskeleton (Shah et al. 2023; Yerima et al. 2023) (Shah et al. 2023; Piccus and Brayson 2020; Rothballer and Kutay 2013). The LINC complex is composed of KASH-domain proteins, known as Nesprins, which are embedded in the outer nuclear membrane and bind to various cytoskeletal elements, including actin filaments, microtubules, and intermediate filament proteins (Rajgor and Shanahan 2013; Cartwright and Karakesisoglou 2014; Mellad, Warren, and Shanahan 2011). SUN-domain proteins, localized to the inner nuclear membrane (Lombardi et al. 2011; Tapley and Starr 2013), form trimers that interact covalently with Nesprins within the perinuclear space (Sosa et al. 2012), and bind indirectly to chromatin and the nuclear lamina (Sosa et al. 2012; Tapley and Starr 2013; Razafsky, Wirtz, and Hodzic 2014). Among the many functions of the LINC complex is its facilitation of nuclear mechanotransduction (Uzer, Rubin, and Rubin 2016; Bouzid et al. 2019; King 2023; Jahed and Mofrad 2019) and control of nuclear localization in muscle fibers (De Silva et al. 2023; Rothballer and Kutay 2013). In addition, the LINC complex mediates chromatin organization and its binding to the nuclear envelope (Shah et al. 2023; Wagh et al. 2021; Piccus and Brayson 2020; Wong, Loo, and Stewart 2021). Genetic disruption of the LINC complex results in aberrant muscle function in a variety of species, including *Drosophila*, *C. elegans*, mice and humans (Lorber, Rotkopf, and Volk 2020; Méjat and Misteli 2010; Cartwright and Karakesisoglou 2014; Auld and Folker 2016; Zhou et al. 2018; Elhanany-Tamir et al. 2012). In *Drosophila,* LINC mutant muscles exhibited altered chromatin organization and weaker association of chromatin with the nuclear periphery (Amiad Pavlov et al. 2023). This is accompanied by enriched epigenetic repressive marks H3K9me3 and H3K27me3, and enhanced binding of the transcriptional repressor Polycomb to a subset of genes. Moreover, LINC mutant muscle cells display reduced levels of the active epigenetic mark H3K9ac and diminished RNAPII binding at specific genomic loci. A theoretical model proposed that reduced chromatin tethering to the nuclear envelope allows for increased chromatin self-attraction, promoting the formation of larger repressive chromatin clusters (Amiad Pavlov et al. 2023). However, the mechanism underlying the reduced RNAPII association with chromatin observed in LINC-deficient muscles remains to be elucidated.

RNAPII operates within dynamic chromatin clusters (Wei et al. 2020; Rippe and Papantonis 2022; Imada et al. 2021), which also contain transcription factors (Trojanowskia and Rippe 2022; W. K. Cho et al. 2018) and regulatory elements (Pancholi et al. 2021; Zhang et al. 2023). These chromatin condensates exhibit a reciprocal regulatory relationship as demonstrated in *Drosophila* (Tullius et al. 2024; Salari, Fourel, and Jost 2024), and are evolutionarily conserved (Zhang et al. 2023; Banigan et al. 2023; Javed et al. 2022; Shao et al. 2022). RNAPII is also involved in multiple stages of transcription and plays a pivotal role in shaping chromatin architecture (Zhang et al. 2023; Chowdhary et al. 2025; Fursova and Larson 2024). Indeed, inhibition of RNAPII leads to global alterations in chromatin organization (Haaf and Ward 1996) and to modulation of nucleosome positioning (Tullius et al. 2024; Gilchrist et al. 2010). Furthermore, RNAPII contributes to chromatin organization during mitotic exit (Zhang et al. 2021), facilitates chromatin loop formation at active gene loci (Banigan et al. 2023; Zhang et al. 2021), and reduces chromatin mobility during active transcription (Nagashima et al. 2019). Thus, RNAPII plays a significant role in nuclear organization beyond its primary transcriptional function.

In this study, we examined correlations in the spatial organization of chromatin and RNAPII within chromatin clusters in the nuclei of live *Drosophila* larval muscles, in wild-type and in LINC mutants, and characterized their relative concentration profiles at the micrometer scale. We demonstrate that in LINC complex mutants, as well as in BAF knocked down nuclei, chromatin cluster size in the nucleoplasm increases, whereas the RNAPII signal decreases within such clusters. Consistently, computer simulations of chromatin clustering as a function of chromatin tethering to the nuclear envelope recapitulate this inverse relationship and explicitly demonstrate that reducing the extent of chromatin tethering to the lamina leads to increased chromatin aggregation in the nucleoplasm and the formation of larger clusters. In addition, we identified asymmetry in RNAPII distribution that correlated with cluster proximity to the nuclear envelope. This asymmetry was reduced in nuclei lacking a functional LINC complex.

Together, our analysis indicates that chromatin tethering to the lamina, mediated by cytoskeletal-nucleoskeletal connections, is essential for limiting chromatin clustering in the nucleus and thereby facilitating its interaction with RNAPII for proper transcription regulation.

## Results

### Chromatin clusters increase in size and show decreased association with RNAPII in LINC mutant muscles

To study the spatial distribution and interactions of chromatin with RNAPII, we imaged live 3^rd^ instar larvae using our custom-designed experimental setup (Lorber, Rotkopf, and Volk 2020). Z-stack images spanning the complete depth of individual muscle nuclei were acquired to generate comprehensive three-dimensional reconstructions. We analyzed wild-type (WT) flies and mutant flies carrying a deletion of the sole *Drosophila SUN*-domain protein, encoded by *koi* gene (*SUN/koi*), both expressing the fluorescent markers, His2B-RFP and Rpb3-GFP (C.-Y. Cho et al. 2022) (see also Materials & Methods).

Fig. 1 shows representative optical cross-sections taken through the centers of wild-type (Fig. 1 WT; panels A and B) and in SUN/*koi* mutant myonuclei in larval muscles. In both genotypes, chromatin is primarily localized at the nuclear periphery, consistent with our previous findings (Amiad-Pavlov, Lorber et al. 2021). Notably, the chromatin layer appears thicker in the SUN/*koi* mutant nuclei (Amiad Pavlov et al. 2023). Remarkably, chromatin is organized into small clusters, as previously demonstrated (Nozaki et al. 2017; Kant et al. 2024; Gelléri, Márton et al. 2024; Miron et al. 2020; Marion Cremer et al. 2017; Y. Li et al. 2022), where RNAPII is closely associated with these clusters, predominately localized to their outer surfaces. Our analysis indicated that repressed chromatin often coincides with the center of chromatin clusters (Supplementary Figure 1). Thus, it is suggested that the His2B-labeled clusters consist of peripheral euchromatin and centrally localized heterochromatin.

**Fig. 1:**
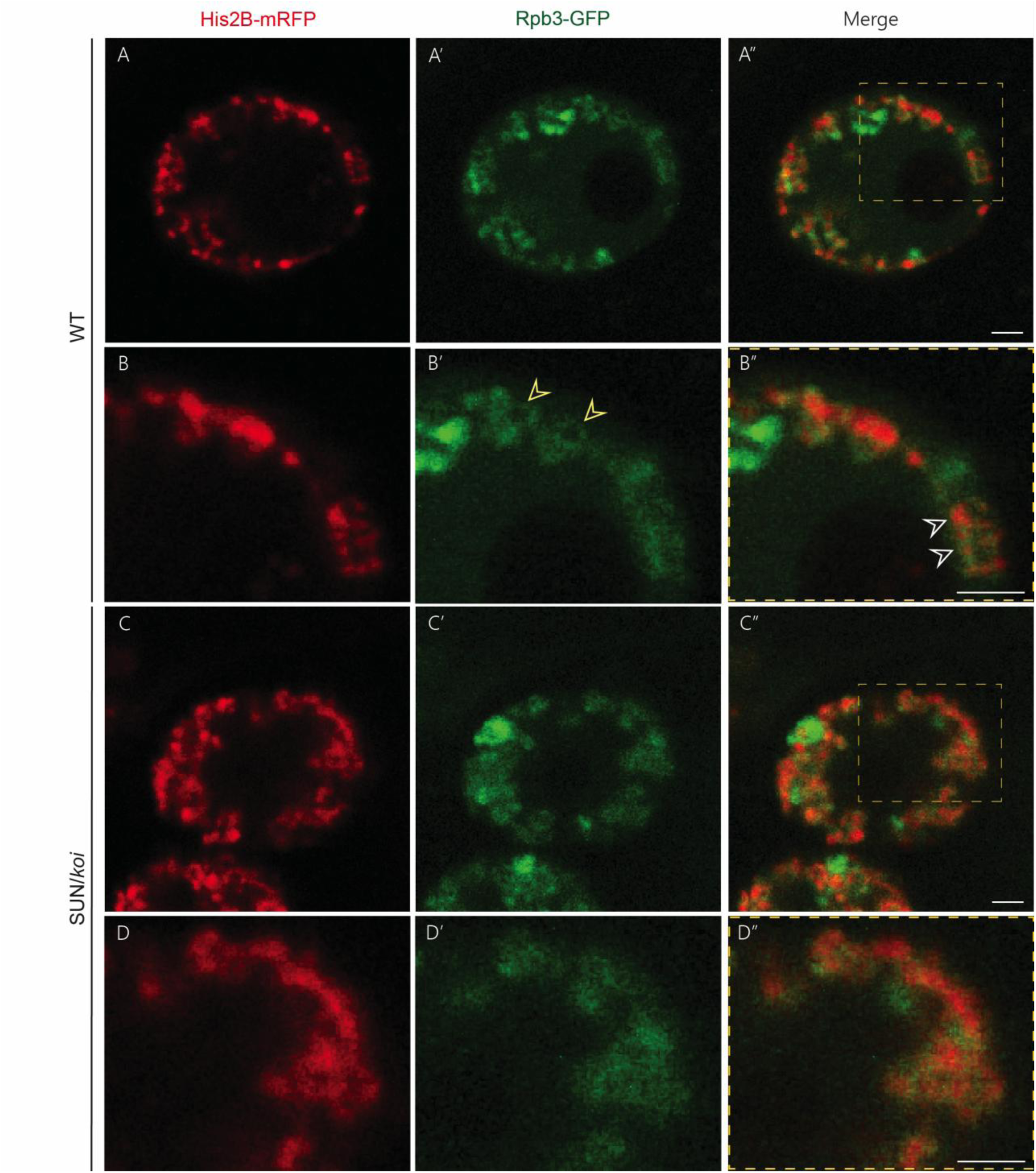
Relative RNAPII-chromatin distribution in control versus SUN/koi mutant muscle nuclei. A-D’’: Single optical XY cross sections taken from WT (A-B’’), or SUN/koi (C-D’’) mutant myonuclei of live, intact *Drosophila* larvae. RNAPII (green, A’, B’, C’, D’), is labeled with Rpb3-GFP and chromatin (red, A, B, C, D), is labeled with His2B-mRFP. Their merged images (A’’, B’’, C’’, D’’) are presented. Marked areas in A’’, and C’’ are enlarged in B-B’’, and D-D’’ respectively. RNAPII surrounds chromatin clusters (yellow arrows) and partially overlaps them (white arrows). In SUN/*koi* mutant nuclei chromatin layer is wider and more diffused and RNAPII levels decrease. (For quantifications see below). Scale bar=2 μm.

Evaluation and quantification of the distributions of chromatin and RNAPII with respect to their reciprocal spatial organization were carried out for each of the chromatin cluster using the following pipeline: First, Local intensity maxima within the imaging dataset were identified using the blob finder tool (using arivis Vision4D software) as seed points, followed by three-dimensional segmentation to identify the location of the cluster’s center of mass (COM) and estimate clusters volumes. No image enhancement or preprocessing of the detected signal were applied prior to this analysis. Subsequently, intensity profiles of His2B-mRFP and Rpb3-EGFP were generated in eight directions, spaced by 45 degrees apart, extending from the COM of each chromatin cluster towards the nucleoplasm, up to a distance of 2 μm, aligned with the x-y axes of the image (Fig. 2A). The fluorescent intensity of each pixel along the profile was calculated by averaging a 3×3 pixel matrix centered on that pixel. Based on these intensity profiles (Fig. 2B, B’, C), we defined a set of quantitative parameters to characterize the relative spatial distributions of chromatin and RNAPII. Note that not all intensity profiles were included in the analysis. For example, in cases where RNAPII signal was located between neighboring chromatin clusters e.g. see Fig. 2C, it was not possible to attribute the signal unambiguously to a specific cluster, consequently, four profiles, two chromatin and two RNAPII, were excluded from the dataset. In addition, cases in which the fluorescent signal did not reach background levels, (defined as a minimal value), within a distance of 2 μm from the cluster COM, were similarly excluded. Additional criteria for profile elimination are detailed in *Materials and Methods.* In total, we analyzed 28 WT myonuclei (YW; Rpb3GFP; His2BRFP) from 11 larvae which yielding 4479 valid profiles pairs, and 30 SUN/*koi* mutant nuclei from 13 larvae, resulting in 3950 valid profiles pairs.

**Fig. 2:**
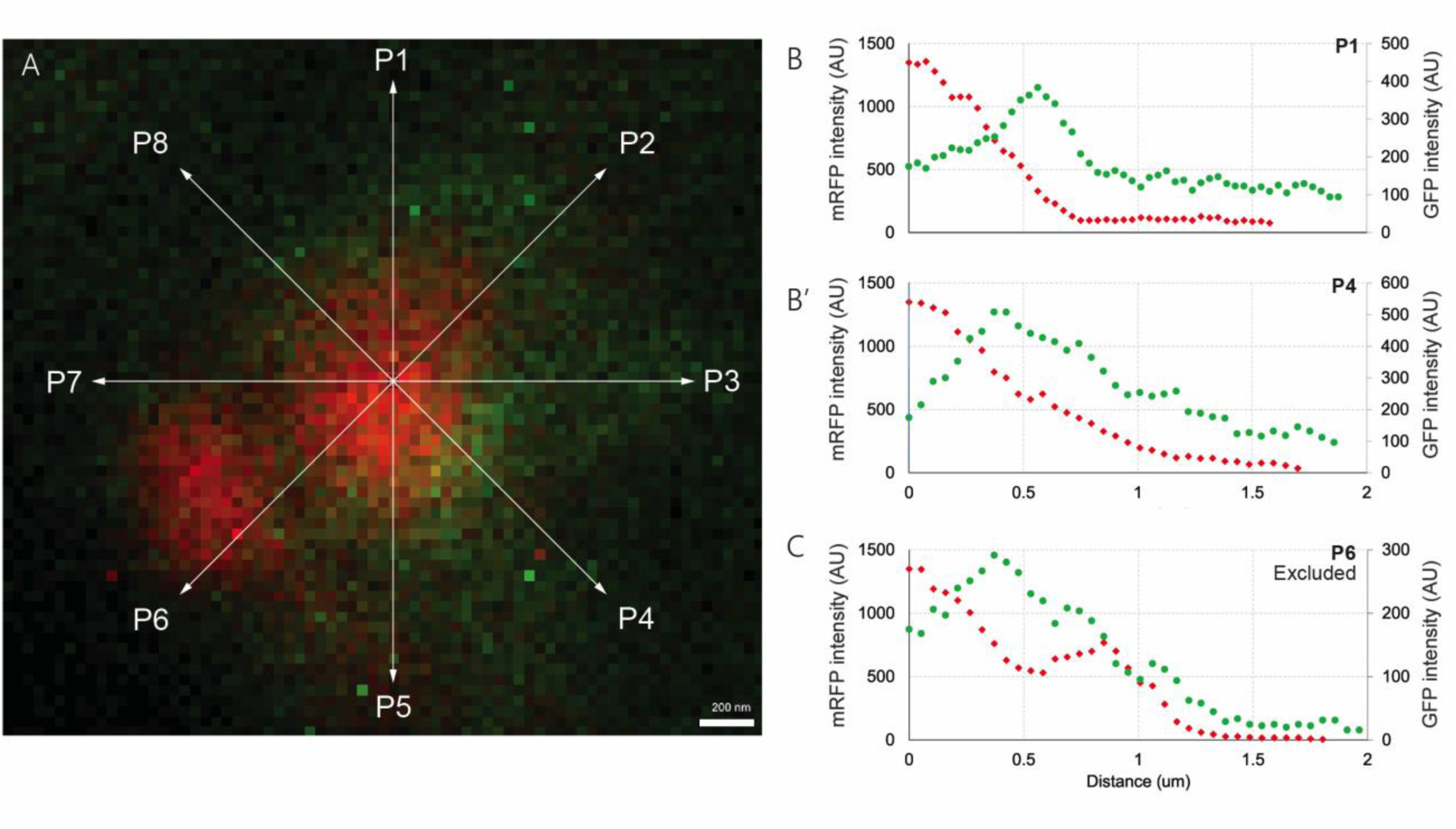
Quantification of chromatin cluster and RNAPII distributions from imaging data. A: Intensity profiles of His2B-mRFP (red) and Rpb3-GFP (green), were generated in eight directions (P1-P8) at the X-Y plan, extending from chromatin cluster’s COM up to 2 µm. B-C: intensity profiles of P1 (B) P4 (B’), and P6 (C), shown in (A). The validity of each chromatin-RNAPII profiles pair was confirmed, (based on a set of criteria, detailed in M&M). (C) Profiles crossing adjacent clusters e.g. P6 were eliminated for both His2B-mRFP and Rpb3-GFP. A total of 4479 valid pairs of profiles from 11 WT larvae and 28 nuclei were sampled. Similarly, 3950 valid pairs of profiles from 13 SUN/koi mutant larvae and 30 nuclei were included.

To quantify the radius of the chromatin clusters we used the “Full Width at Half Maximum” (FWHM) estimation (de Coninck et al. 2018; Georgiev et al. 2015). For each profile we calculate half the difference between the maximum intensity and the first local minimum, then determined the x-position corresponding to this half-maximum intensity (interpolated when necessary; Fig. 3A). On average, the radius of a chromatin cluster in *SUN/koi* mutant larvae was larger than that of WT: 366.8 nm versus 316.9 nm respectively, p<0.0001). Assuming chromatin clusters are perfect spheres, the cross-sectional area of the *SUN/koi* clusters would be 34% greater than that of WT clusters, and their volume 55% larger. Additionally, the location of maximal intensity was found to be 38.3 nm from the cluster COM in WT clusters, and 44.3 nm in *SUN/koi* myonuclei (p<0.0001) (Fig. 3B,C). These results imply that chromatin organization and packaging differ significantly between *SUN/koi* mutant nuclei and WT. Beyond the size difference, chromatin clusters in *SUN/koi* mutants also exhibited larger standard deviation in size based on a Wilcoxon rank test: 143 nm compared to 125 nm (*p* < 0.001).

**Fig. 3:**
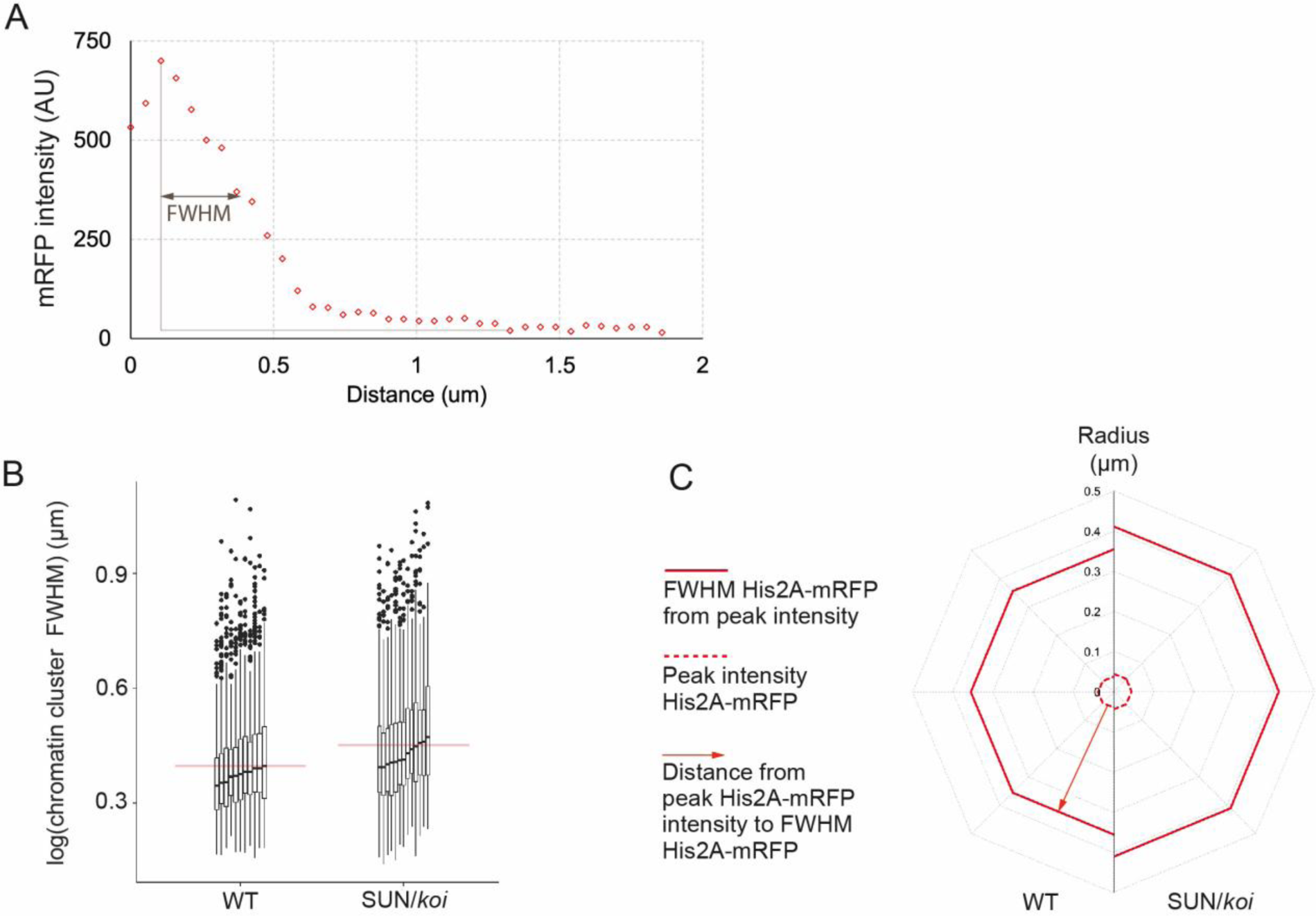
Chromatin clusters are enlarged in SUN/*koi* mutant nuclei compared to controls. (A) Quantification of chromatin cluster size was performed by calculating the Full Width at Half Maximum (FWHM) for each cluster radius, defined as the distance corresponding to half the difference between the maximal and minimal His2B-mRFP (red) intensity values. (B) Summary of the FWHM values for all clusters, with each vertical line representing an individual larva. SUN/koi mutants show significantly larger FWHM values compared to WT (366.8 nm vs. 316.9 nm, p-value=2.3·10^-5^) and exhibit a larger standard deviation (143 nm vs. 125 nm, p-value<0.001). The horizontal red line indicates the population average. (C) The spatial distribution of chromatin in SUN/*koi* mutants (right half circle) compared with WT (left half circle). Red line represents FWHM range, which is greater in the mutant. In addition, the distance between the cluster COM and peak intensity is greater in SUN/koi mutants (44.3 nm) compared to WT (38.3 nm), p-value=5.1·10^-5^), with higher standard deviation (73.9 nm vs. 64.5 nm, p-value=0.0072).

A similar approach was used to characterize RNAPII distribution. For each chromatin cluster we measured the FWHM of RNAPII from its peak intensity in two directions: toward the nucleoplasm and towards the cluster’s COM (Fig. 4A). These measurements revealed a significantly reduced distribution range of RNAPII in SUN/*koi* mutants compared to WT. In WT nuclei, RNAPII extended 320.3 nm toward the nucleoplasm and spanned a total width of 383.3 nm, whereas in SUN/*koi* mutants, the extension toward the nucleoplasm was 279.6 nm with a total span of 340.2 nm (*p* = 0.007, and *p* = 0.01, respectively, Fig. 4B, B’). Furthermore, the standard deviation in RNAPII distribution, both toward the nucleoplasm and in total width, was significantly greater in SUN/*koi* mutants (*p*= 0.005, and *p* = 0.0002, respectively), consistent with the increased standard deviation observed in the chromatin radius. Taken together, our measurements indicate that LINC mutant nuclei exhibit a significant increase in chromatin cluster size and a reduced range of RNAPII distribution surrounding each cluster.

**Fig. 4:**
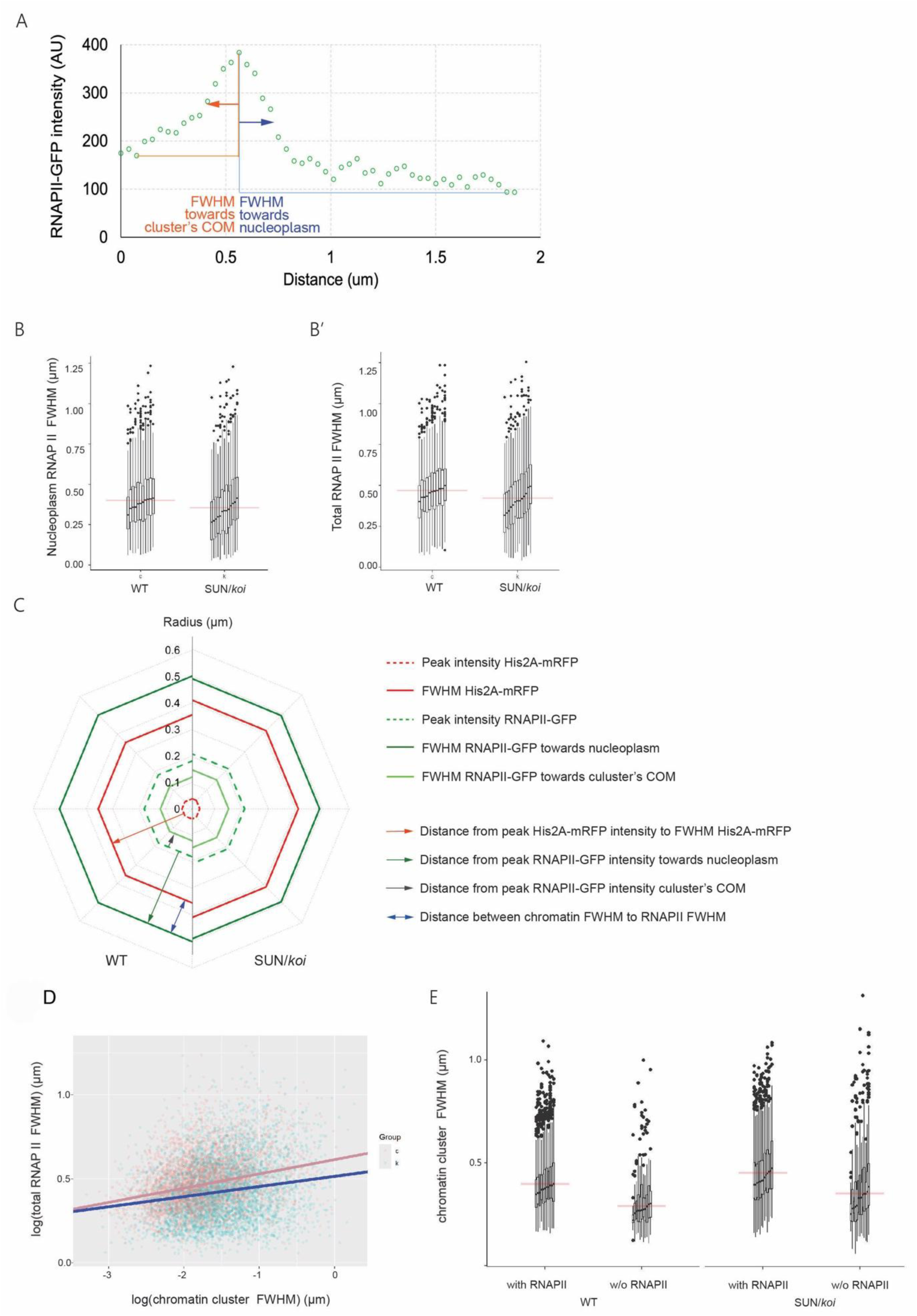
The range of RNAPII signal around chromatin clusters is reduced in SUN/koi mutants compared with WT. Quantification of RNAPII distribution was based on the FWHM of the Rpb3-GFP signal, measured toward the nucleoplasm and toward the cluster COM, corresponding to the sum of the red and blue arrows. (B) FWHM values of RNAPII distribution toward the nucleoplasm per chromatin cluster are significantly lower in SUN/koi nuclei (279.6 nm) compared to WT controls (320.3 nm; *p*=0.007) and exhibit higher standard deviation (181 nm vs. 166 nm, *p*=0.005). (B’) Total RNAPII distribution, calculated as the sum of FWHM values, toward the nucleoplasm and the cluster COM, is reduced in *SUN/koi* mutants (340.2 nm) compared to WT controls (383.3 nm, *p* = 0.01) and shows higher standard deviation in the mutant (176 nm vs. 159 nm, *p =* 0.0002). (C) Spatial distribution of chromatin and RNAPII in WT (left) and SUN/*koi* mutants (right) myonuclei, showing the distance between chromatin FWHM (red line) and RNAPII FWHM toward the nucleoplasm (green line). This distance (blue double arrowheads line) is significantly shorter in SUN/koi mutants compared to WT control (104.5 nm, vs. 163.6 nm *p* = 0.002). (D) Correlation between RNAPII and chromatin distributions. The graph shows FWHM values of the RNAPII layer plotted against FWHM values of chromatin clusters in SUN/*koi* mutants (blue dots), and in WT controls (orange dots). In WT nuclei, RNAPII layer width (FWHM) is strongly positively correlated with chromatin cluster radius (FWHM) (*p*<2 x 10^-16^). This relationship is weaker in SUN/koi mutant nuclei, where RNAPII layer width is less sensitive to cluster size (*p* = 0.009). (E) Comparison between chromatin cluster size with, or without RNAPII. In WT nuclei, the size of chromatin clusters (FWHM) lacking RNAPII were significantly smaller than those associated with RNAPII (223.7 nm vs. 317.5 nm, p = 3.4 x 10^-69^). In SUN/koi mutant myonuclei, clusters lacking RNAPII were also smaller but larger than their WT counterparts (276.1 nm vs. 367.6 nm; *p* = 9.42×10^-92^).

An additional parameter used to characterize the spatial relationship between RNAPII, and chromatin was the distance between FWHM of chromatin and that of RNAPII, measured along the profiles toward the nucleoplasm (Fig. 4C). On average, this distance was significantly shorter in *SUN/koi* mutants compared to WT (104.5 relative to 163.6 nm; *p* = 0.002), consistent with the reduced RNAPII distribution and enlarged chromatin cluster size observed in the mutants.

Next, we examined the relationship between chromatin cluster size and the distribution of the associated RNAPII. To this end, we plotted the total RNAPII width (the sum of both directions) against the radius of its corresponding chromatin cluster for all intensity profiles pairs (Fig. 4D). In both WT and *SUN/koi* mutants, RNAPII distribution correlated with chromatin cluster radius; however, this correlation was notably weaker in the SUN/koi mutant (*p* < 2e-16, for WT; *p* = 0.009 for *SUN/koi*). This finding is consistent with the reduced RNAPII association with chromatin in *SUN/koi* mutants and suggests diminished sensitivity of RNAPII distribution to the chromatin cluster size.

We also investigated whether chromatin cluster size is influenced by low RNAPII presence in its vicinity. To address this, we analyzed clusters with at least four RNAPII profiles (out of eight) that were excluded from the initial analysis due to their total low intensity (see materials and methods for details on exclusion criteria). Chromatin clusters associated with low RNAPII levels were significantly smaller in both WT and SUN/*koi* mutants. Nevertheless, the relative difference in cluster dimensions between WT and mutants remained consistent with prior analyses (Fig 4E). In WT nuclei, the chromatin clusters lacking associated RNAPII were smaller than those associated with RNAPII (radius=223.7 nm relative to 317.5 nm; Fig. 4E left panel). Similarly, in SUN/*koi* mutant nuclei, clusters with low RNAPII levels were smaller (radius=276.1 nm, relative to 367.6 nm; Fig. 4E, Right panel). These differences were statistically significant in both cases (*p* = 3.4×10^-69^, and *p* = 9.4x 10^-92^, respectively).

In summary, our findings suggest two important differential features of chromatin microscale organization in SUN/koi mutants: enlarged chromatin cluster size and reduced RNAPII levels per chromatin cluster. The enlarged size of chromatin clusters in *SUN/koi* mutants, reflects an intrinsic feature of their spatial chromatin organization independent of RNAPII presence.

To further address the link between binding of chromatin to the nuclear envelope and the size of chromatin clusters, we knocked down Barrier to Autointegration Factor (BAF) in larval muscles using a muscle specific driver combined with BAF RNAi (BAF-KD). BAF, a DNA cross-linker protein, binds to DNA and to LEM-domain proteins located at the nuclear envelope, independent of the LINC complex (Andrés and González 2009; Sears and Roux 2020; Barton, Soshnev, and Geyer 2015; Gruenbaum and Foisner 2015; Marcelot et al. 2021; Sobo et al. 2024). In BAF-KD muscles, the chromatin layer is relatively dispersed (Fig. 5B) compared to WT (Fig. 5A) and chromatin cluster size is significantly enlarged (453.3 nm, 316.9 nm, respectively, *p*=4.5X10^-9^). Similar to the trend found in the LINC mutants, lack of RNAPII in BAF RNAi nuclei correlated with reduced chromatin cluster size to 310.4 nm (*p*=4.2×10^-10^, Fig. 5D).

**Fig. 5:**
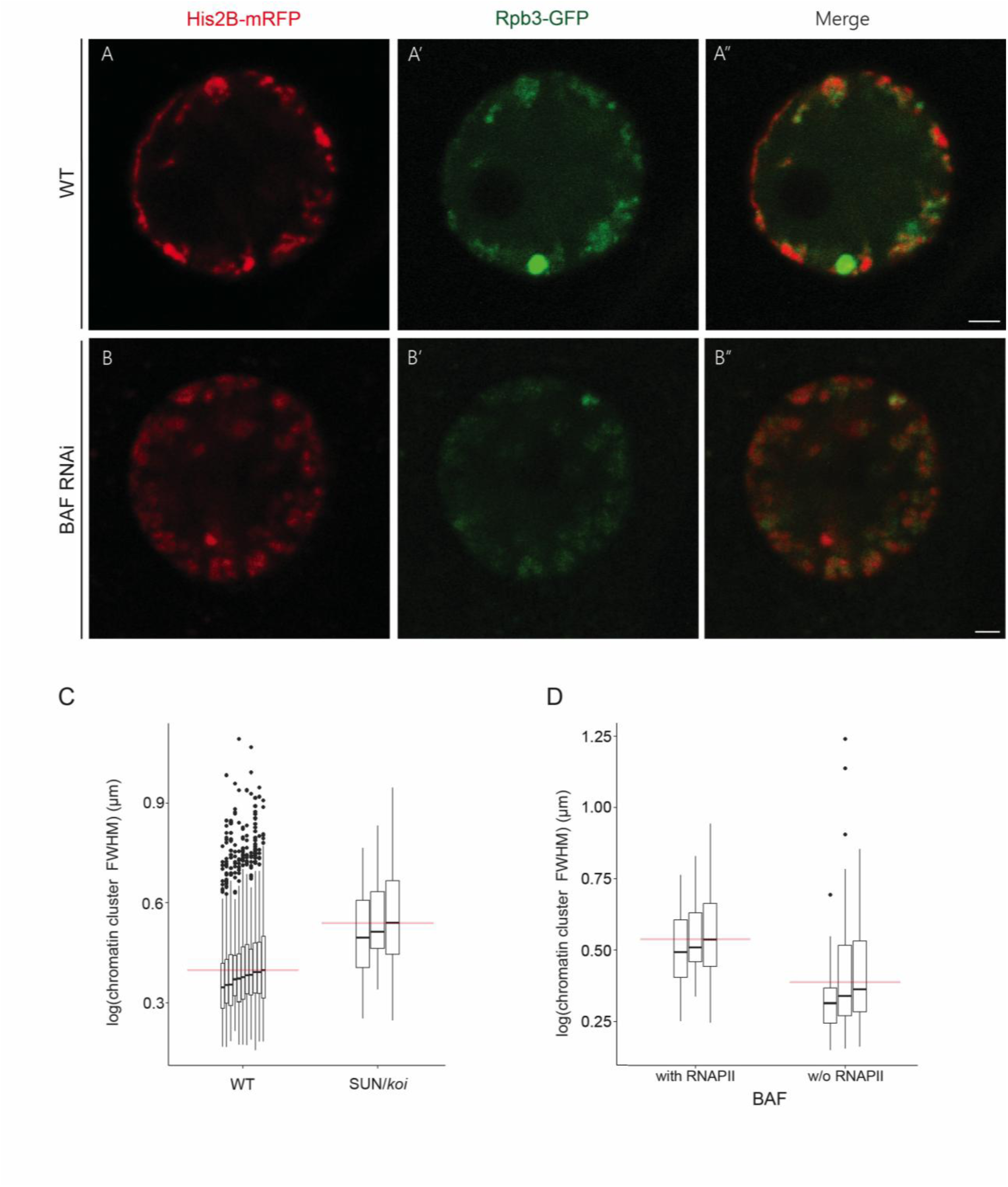
Chromatin and RNAPII organization in BAF RNAi nuclei. WT (A-A’) and BAF knockdown (B-B’) myonuclei labeled for His2B-mRFP (A,B) and Rpb3-GFP (À, B’) and their merged images (A’, B’) of myonuclei from live, intact larva. Note that in BAF RNAi nuclei chromatin clusters shift from the nuclear periphery (C). Chromatin clusters in BAF RNAi nuclei are significantly larger than WT ones: 453.3 nm compared to 316.9 nm, (*p*=4.5X10^-9^). (D) In the absence of RNAPII in BAF-KD nuclei, chromatin cluster FWHM reduces (310.4 nm, *p*=4.2×10^-10^).

These results support the notion that reduced binding of chromatin to the nuclear envelope rearranges chromatin organization resulting with larger cluster sizes.

### Computer simulations demonstrate inverse relations between reduced chromatin binding to the nuclear envelope and increased chromatin cluster size

Previously, we showed both experimentally and theoretically that reduced chromatin attachment to the nuclear envelope following overexpression of lamin A/C correlates with the collapse of peripheral chromatin into the nuclear center (Amiad Pavlov, Lorber et al., 2021; Amiad Pavlov et al., 2023). Here, we show that when we use computer simulations to model reduced chromatin binding to the lamina (relevant to the experiments on the LINC mutant and BAF KD), chromatin clusters increase in size. This demonstrates how both the heterochromatin and euchromatin are affected by reduced chromatin binding to the nuclear lamina in the LINC mutants. The simulations are a guide to appropriate parameter ranges that recapitulate both the reduction in the effective binding of LADs to the lamina in the LINC mutants and chromatin self-interactions, as well as chromatin interactions with RNAPII (Adame-Arana et al., 2023; Amiad Pavlov, Lorber et al., 2021).

The simulations model one chromosome as a multiblock copolymer with three types of repeating blocks: active (type A, euchromatin), inactive (type B, heterochromatin), and LAD (type C, heterochromatin that binds to the lamina). Additional details of the simulation model are presented in Materials and Methods (M&M) and Appendix A. First, irrespective of chromatin type, the attractive interaction between the chromatin monomers leads to the collapse of the polymer, causing phase separation from the solvent due to the self-attraction of the chromatin modeled as beads on a chain (De Gennes 1979, Rubinstein, Colby 2003, Kumar et al., 2019). Furthermore, the strong interaction between inactive monomers (as a model of heterochromatin) (Fig. 6 blue, BB, monomers) compared to the relatively weaker interaction between active monomers (as a model of euchromatin) (Fig 6. red, AA monomers) causes microphase separation of the active and inactive blocks with a smaller reduction of the system free energy compared to that induced by chromatin-nucleoplasm interactions (for bead explanation, see Appendix A). Consequently, the inactive monomers from the same and neighboring blocks collapse to form an inactive core surrounded by a shell of active monomers. Topological constraints due to chain connectivity and self-avoidance of the chain, as well as the attractive interaction energy gain, play the major role in the formation of the “strings of micelles” structures seen in Supplement Figure 2 (Adame-Arana et al., 2023). In addition, another set of beads mimicking RNAP II attaches to the active blocks due to attractive and binding interactions. This results in effective shielding of the attractive interaction of active chromatin blocks (AA) from each other, eventually leading to their dilution in the nucleoplasm. This modifies the string of micelles assemblies so that the A blocks are bound to and sometimes coated with RNAP II. In addition to these effects, the LAD chromatin beads (C) strongly associate with an additional set of confining beads that represent the nuclear envelope. We vary the LAD (C) fraction in the chain to mimic the wild type, as well as variants like the LINC mutant, where the binding of the periodically distributed LAD (C) blocks to the confining beads is reduced. To correlate (but not match precisely) with *Drosophila* nucleotide sequence data, we keep the number of LAD blocks m = 100. The total number of monomers in the linear polymer, *N = n (N_A_+N_B_) + mN_C_*, where *n* is the number of active and inactive blocks and m is the number of LAD blocks. We adopt a polymer of size *N* = 50000, where the fraction of different types of monomers and block lengths is tuned. We fix the number of A and B blocks to be identical with *n* = 12, and depending on the LAD fraction, *N_A_*, *N*_B_, and *N*_C_ can then be determined. For the largest LAD fraction we studied, (*m Nc)/N = 1/2*, we observe peripheral localization of all the chromatin domains, with about half of the LAD monomers adhered to the confinement wall. Furthermore, the topological constraint of a connected chain effectively pulls the micelles (consisting of A, B, and RNAP II beads) towards the LAD section of the chromatin and hence close to the confining wall (modeling the lamina), leading to peripheral organization of chromatin domains. We mimic the reduced binding of the LAD (C) beads and the wall, as a way of representing the LINC mutants. To do this, we effectively eliminate a certain fraction of the LAD (C) beads and “convert” them to B-type heterochromatin beads. This conserves their post-translational modifications (e.g., methylation) and their identity heterochromatin. Thus, *N*_C_ decreases while *N*_B_ increases, conserving the total number of heterochromatic (C and B types) polymer beads. This effectively reduces the effective attraction of all the chromatin to the confining wall (lamina); thus, the A, B, RNAP II micelles are pulled less strongly towards the confining wall. These micellar assemblies thus tend to diffuse towards the center of the nucleus since delocalization increases their entropy. In Fig. 6, the LAD (C) fraction is reduced as one goes from the leftmost panel to the right. At a LAD (C) fraction of 45%, we already see dissociation of micelles from the nuclear periphery, and the overall chromatin concentration is more dilute. A further reduction in the LAD fraction accentuates this effect, leading to further dissociation of chromatin from the spherical wall and an even more homogeneous distribution of micelles in bulk.

**Fig 6:**
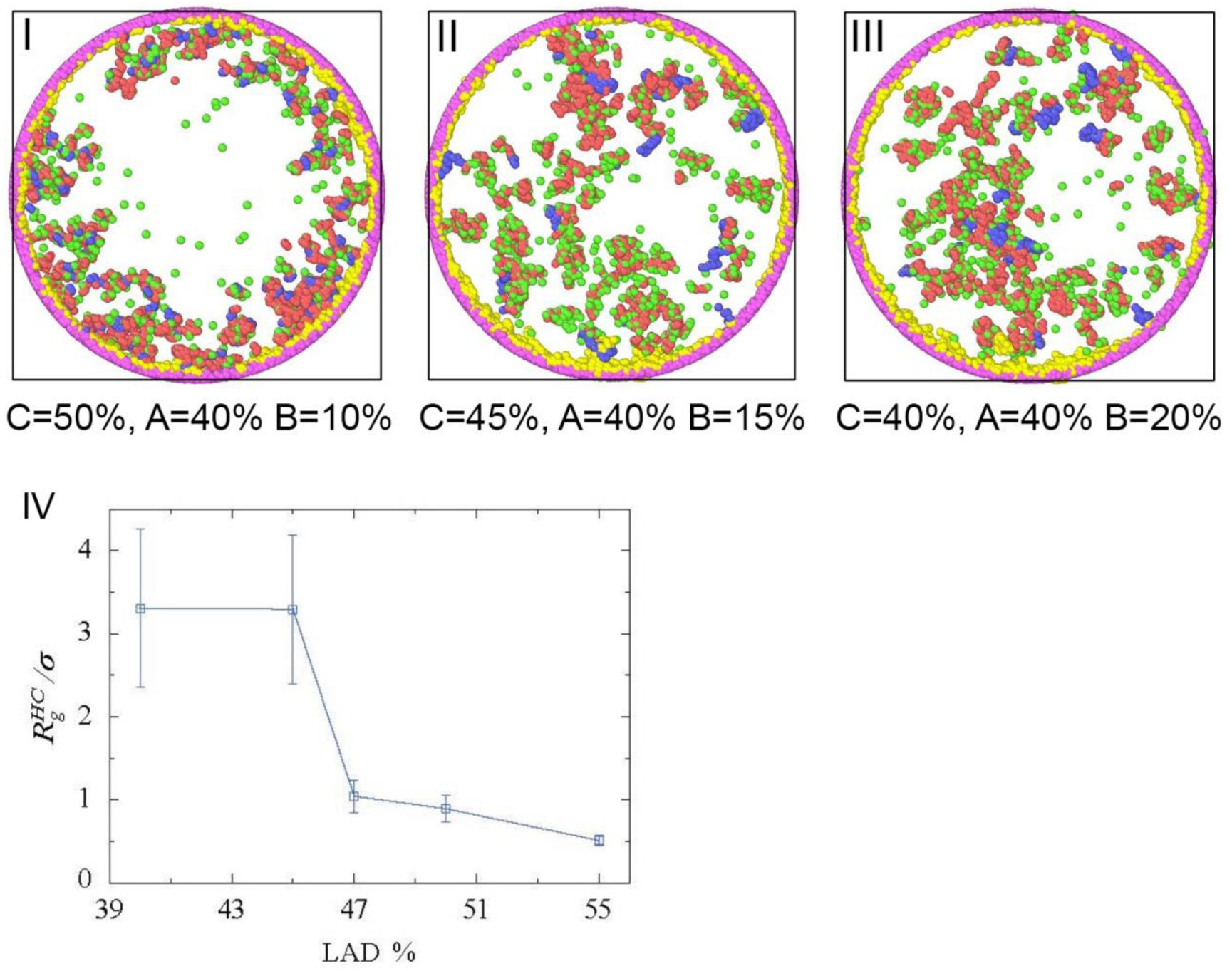
Simulation of the spatial organization of chromatin domains and RNAP II within nuclear confinement. Panels I, II, and III show the configurations of chromatin domains (A-type red, B-type blue, C-type yellow, and RNAPII, green) for different fractions of each chromatin type. From left to right, the LAD fraction (C) is decreased, and the B monomer fraction increases. The active A monomer fraction is fixed at 40% of the total monomers of the chain. The spherical beads shown in pink represent the confinement beads (which model the nuclear envelope). (I) LAD fraction fixed at 50%, and inactive chromatin B at 10%, showing peripheral organization of all the chromatin domains. The A and B chromatin domains show a micelle-like organization consisting of an inactive (B) core and an active (A) shell. The active shell consists of active monomers that are covered (wet) by the green beads (RNAPII) due to bonded and non-bonded attractive interactions between them. (II) and (III) show LAD fractions reduced to 45% and 40%, respectively. Both show a significant reduction in the peripheral organization of chromatin domains. The volume fraction of the total number of chromatin beads in the system is globally fixed at 0.05, while the fraction of RNAPII beads in the system is 0.008. (IV) Radius of gyration of inactive (B) cluster core. The radius of gyration of the inactive (B) cluster core (y-axis) is plotted as a function of the LAD fraction (x-axis) of total monomers of the chain. The cluster size is measured in units of the monomer diameter σ. The apparent decrease in the cluster size is observed with an increase in LAD fraction.

In addition to a change from strongly peripheral to very weakly peripheral organization that occurs in the simulations as we reduce the LAD (C) fraction (modeling the LINC mutant), another important change takes place: the size of the inactive (heterochromatin) cluster core increases sharply as shown in Fig. 6D. The mean radius of gyration of inactive cluster core of micelles (as computed over equilibrated configurations) increases as the LAD fraction decreases. This occurs due to the increasing content of inactive (B) monomers as the LAD (C) are converted to (B) to mimic the LINC mutants, where the binding to the nuclear envelope is reduced. To conclude, both effects, a change from strongly to very weakly peripheral organization and an increase in the cluster size of the inactive (B) beads, are due to the decrease in the LAD fraction mimicking the effect of LINC mutations, which reduces the binding of the chromatin to the nuclear envelope.

In conclusion, the simulation results demonstrate a mechanistic link between the reduced LAD fraction (observed in the LINC mutant) and increased chromatin cluster size, consistent with the experimental data.

### Asymmetric arrangement of RNAPII and chromatin clusters correlates with nuclear envelope proximity

Current models of chromatin organization propose that repressed heterochromatin is typically associated with the nuclear lamina, whereas active euchromatin is displaced toward the nucleoplasm (Oji et al. 2024; Sood and Misteli 2022; Haws et al. 2022). To investigate chromatin and RNAPII spatial arrangement in live muscle nuclei, we analyzed their distribution as a function of their distance from the nuclear envelope, as well as the direction of their surface area, in both wild type and SUN/*koi* mutant nuclei.

To this end, chromatin clusters whose COM was located in similar plane as the nuclear COM were selected (Fig. 7A). The fluorescent profiles of the clusters that aligned with the radius of nucleus were further analyzed, and their parameters were plotted as a function of their proximity to the nuclear envelope and surface direction (Fig. 7 B,C). The intensity profiles facing the nuclear COM were labeled as IN, and conversely, the profiles oriented towards the nuclear envelope were denoted OUT. In total, 131 clusters from WT nuclei and 141 from *SUN/koi* mutant were analyzed.

**Figure 7:**
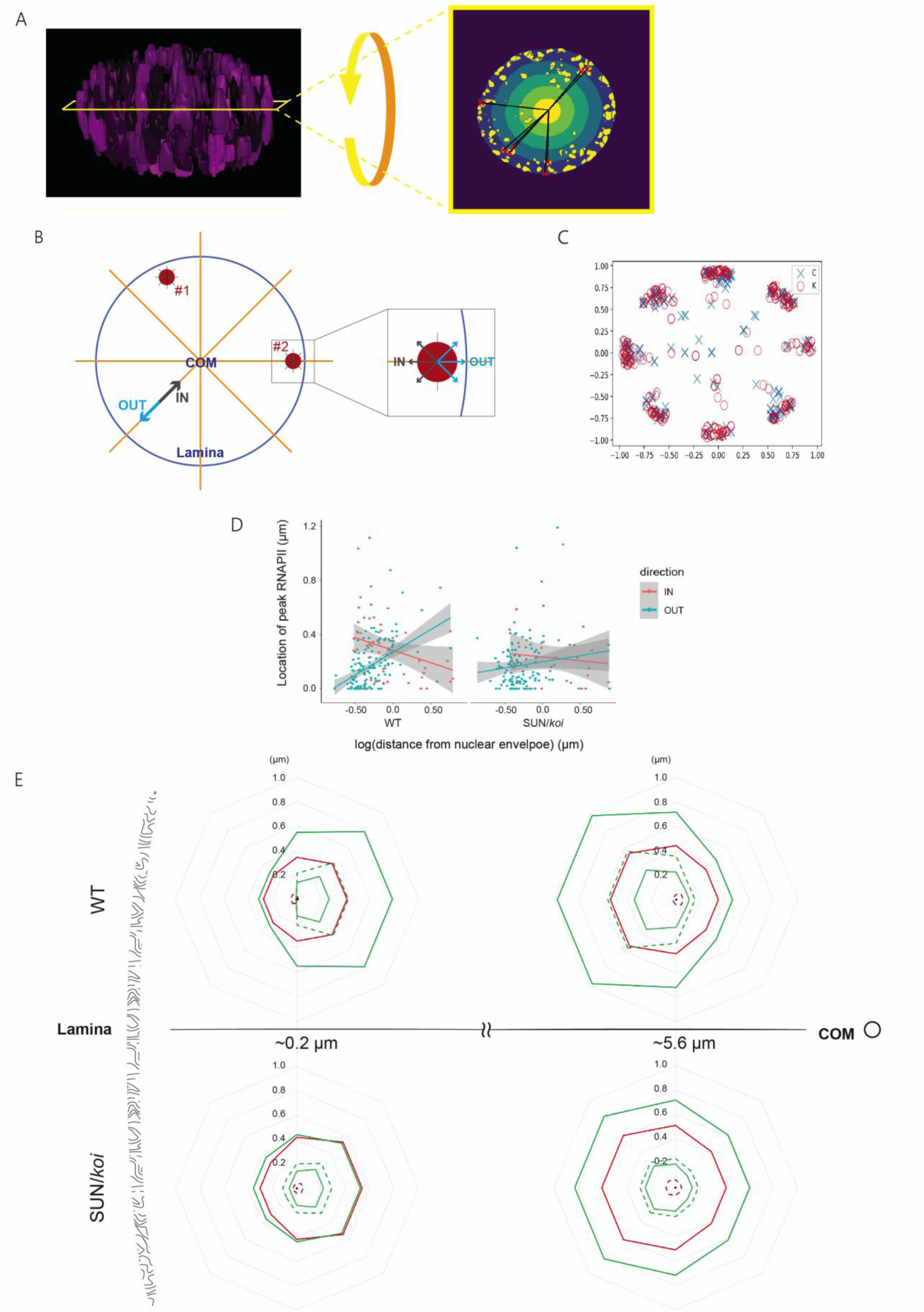
The spatial distribution of chromatin and RNAPII as a function of cluster proximity to the nuclear envelope. (A and B): Selection of clusters for chromatin and RNAPII orientation analysis. Only clusters whose COM aligned with the nuclear COM in the z-direction were included. Furthermore, the analysis included only clusters having profiles directed towards the nuclear COM (B). In this example, none of the profiles of cluster #1 were oriented towards the nuclear COM, and hence, omitted from this analysis. In cluster #2, the profile extending toward the nuclear COM was designated ‘IN’, and the profile extending towards the nuclear envelope was designated “OUT” (C) The population of WT (blue) and mutant SUN/*koi* (red) chromatin clusters included in this analysis shown on a single normalized nuclear cross section. In WT nuclei the selected clusters comprised yielded 57 IN-oriented profiles and 134 OUT-oriented profiles. From the SUN/*koi* mutants clusters we obtained 29 IN-oriented profiles, and 134 OUT-oriented profiles. (D) Graph showing the change in peak RNAPII location relative to cluster COM as a function of cluster distance from nuclear envelope and its surface direction. In the direction facing the nuclear envelope marked OUT, RNAPII peak location is close to the clusters COM and shifts away as the distance from the lamina increases (p-value=0.00017). For the surface facing the nuclear COM, noted IN, peak RNAPII location shows an opposite trend (p-value=0.056), getting closer to the cluster’s COM as its distance from the lamina increases.

The parameters characterizing different aspects of chromatin and RNAPII distributions included: the location of peak chromatin and RNAPII intensities, the width of the RNAPII layer (measured both toward the nucleoplasm, and toward the cluster COM), and chromatin cluster radius from its peak intensity. A representative graph showing the location of peak RNAPII as a function of cluster distance from the nuclear envelope is presented in Fig. 7D, for profiles directed towards the nuclear envelope (blue), or the nuclear center (red). Proximal to the nuclear envelope, on the OUT surface (blue), peak RNAPII intensity is close to the cluster’s COM and gradually shifts outwards for clusters located away from the nuclear envelope (p=0.00017). The location of peak RNAPII intensity on the IN surface (orange) shows an opposite trend, yet weaker, as it gets closer to the cluster’s COM in distal chromatin clusters (p=0.056).

Notably, this spatial asymmetry is significantly reduced in SUN/koi mutants, suggesting a role for the LINC complex in promoting chromatin cluster asymmetry. Graphs summarizing these analyses are presented in Supplement Figure 3.

The analysis in Fig. 7E provides a graphical summary of all measurements, illustrating the distributions of chromatin and RNAPII in WT and *SUN/koi* mutants myonuclei for clusters positioned either proximal to the nuclear envelope (average distance: 0.2µm) or distal (average distance: 5.6 µm). In WT nuclei, chromatin and RNAPII clusters proximal to the nuclear envelope are more dispersed toward the nuclear center than toward the nuclear envelope. This tendency is inverted in clusters distal to the nuclear envelope. Notably, this asymmetry was not observed in the *SUN/koi* mutant nuclei either in clusters proximal or distal to the nuclear envelope. In summary, our measurements show that the organization of RNAPII and chromatin depends on the proximity of chromatin to the nuclear envelope and on its orientation, either facing the envelope or nuclear interior. Notably, this dependence is lost in LINC mutants, suggesting that the LINC complex regulates nuclear envelope-dependent asymmetry of chromatin architecture.

These trends were attenuated in the SUN/*koi* mutant and the difference between them is not significant (OUT and IN p-values are 0.56 and 0.31, respectively). Detailed results of all the tested parameters are found in Appendix A. (E) A scheme summarizing all the parameters analyzed. The chart shows that chromatin and RNAPII in WT exhibit directional organization that depends on the distance of cluster from the nuclear envelope and surface orientation. This asymmetric organization is lost in the SUN/*koi* mutants.

## Materials and Methods

### Fly Stocks and Husbandry

The following stocks were used: *koi^84^*/Cyo-dfd-eYfp (FBst0025105) described previously (Kracklauer et al. 2007) GAL4Mef2.R (FBst0027390), w; EGFP-Rpb3; His2Av-mRFP (C.-Y. Cho et al. 2022); koi^84^ was recombined with EGFP-Rpb3; His2Av-mRFP on 2^nd^ chromsome. BAF-RNAi (PRID:BDSC_36108). All crosses were carried and maintained at 25◦C and raised on cornmeal agar. Homozygous SUN/koi mutant larvae were selected by non-Cyo-YFP and confirmed with mis-localization/aggregation phenotype of muscle nuclei.

UAS-2E12LI-EGFP(III)/TM6B (live H3K27me3-GFP mintbody) obtained from H. Kimura, Tokyo Institute of Technology, Yokohama, Kanagawa 226-8503, Japan).

### Imaging nuclei in a live *Drosophila* larvae

Myonuclei of live 3^rd^ instar larvae were imaged using a custom-designed apparatus (Lorber, Rotkopf, and Volk 2020). In brief, the device features a central plastic bar with a longitudinal groove where the larva is positioned. The larva is glued at its anterior and posterior ends to glass capillaries and encapsulated in hydrogel. For imaging, the assembled device mounted onto a microscope stage. All the images presented in this study were acquired using this device.

#### Microscopy and image acquisition

The imaging protocol was previously described in detail (Amiad-Pavlov, Lorber et al. 2021). Shortly, third instar wandering larvae were collected form vials walls, and placed in tap water for about 4 hours before the experiment to reduce their mobility (which is reversable when re-exposed to air). The larvae were then inserted into the imaging device.

Z-stacks imaging of the live nuclei was performed using an inverted Leica SP8 STED3X microscope, equipped with internal Hybrid (HyD) detectors and Acusto Optical Tunable Filter (Leica microsystems CMS GmbH, Germany) and a white light laser (WLL) excitation laser. A white light laser was used in reflection mode to image the sarcomeres surrounding the nuclei and confirm nuclear identity. RFP emission signal was collected at the range of 597– 699 nm and GFP emission in a range of 488 – 559 nm, based on a spectral analysis performed on the larvae nuclei. All the nuclei were visualized with a HC PL APO 86x/1.20 water STED White objective, NA=1.2, at 12-bit depth. The field of view and laser intensity were adjusted for each sampled nucleus to ensure optimal acquisition quality.

#### Image processing and data analysis

Imaged nuclei were selected from larval body wall muscles randomly. For analysis, we selected only nuclei that did not move during sampling and that their adjacent sarcomeres could be detected in the imaging planes.

Image analysis was performed as follows: Firstly, using Arivis Vision4D, chromatin clusters along the entire Z-axis of the nucleus were detected. Secondly, an in-house pipeline (see below) developed in Python to analyze all clusters as presented in this work.

#### Pipeline for the image analyses

Using Arivis, chromatin clusters were identified using the ‘blob finder’ filter that identifies local maxima, without any prior enhancement or filtering of the raw image. The output of this analysis, i.e., x, y, z of each cluster’s COM together with the raw image data and segmented clusters were used as input for the custom Python code for both channels (chromatin and RNAPII).

A set of conditions was formulated to ensure that the analyses performed only on valid clusters:

1. Valid RNAPII and chromatin profiles are free from missing values.
2. Only intensity profiles of chromatin and RNAPII with a signal RNAPII area within 95% of the population, per nucleus, were used for the various analyses. Profiles included within the lowest 5% were used only for the analysis of chromatin cluster diameter lacking RNAPII.
3. To exclude ambiguous data obtained between two chromatin clusters, valid RFP fluorescent profiles were defined as those that exhibit local maximal intensity up to 5 pixels from the cluster’s COM, and a minimal value which is at least 30% lower than its maximal value. In addition, the intensity should go down without a significant increase along its length.
4. Valid RNAPII profile should decrease by at least 50% from its maximal value along a distance of 2 µm.

The valid extracted intensity profiles were subjected to the additional analyses.

Code was written in Python 3. NumPy library (v. 1.26.4) (Harris et al. 2020) was utilized for arrays manipulation together with pandas library (v. 2.2.2) (The pandas development team 2020) for tables analysis. SciPy.Stats package was utilized for initial statistical analysis, and SciPy.Signal, was utilized for helping in analyzing the intensity profiles (v. 1.13.1) (Virtanen et al. 2020). OpenCV library (v. 4.11.0) (Bradski 2000) was utilized for image analysis together with Scikit-Image library (v. 0.23.2) (van der Walt et al. 2014) and SciPy.Ndimage package. Scikit-Learn library (v. 1.5.0) (Pedregosa et al. 2011) was utilized for data analysis. Matplotlib library (v. 3.6.2) (Hunter 2007) was utilized for visualization and figures preparation.

### Statistical analysis

Variables were compared between control and *SUN/koi* using linear mixed effects models, with genotype as a fixed factor, and larva and nucleus as random factors.

Variables with heavily right-skewed distributions were log-transformed before the analysis.

Reported mean values are extracted from the results of mixed effects models, therefore they may be slightly different between analyses, depending on the groups included in the data for each specific model.

For testing the added effect of distance from the nuclear envelope, distance and genotype (and their interaction) were entered as fixed factors, and larva and nucleus as random factors.

Analyses were run in R, v. 4.5.1, using ‘lme4’ and ‘lmerTest’.

## Discussion

Chromatin organization in differentiated cells requires long-term transcription regulation, yet studies describing the precise chromatin 3D distribution, together with its associated transcription machinery are still missing. Here, the specific 3D chromatin organization and its associated RNAPII distribution were studied in mature muscle tissue using image analysis of live *Drosophila* larvae. The analysis was performed both in wild type and in SUN/*koi* mutants, to reveal the contribution of the LINC complex to the regulation of chromatin organization. We previously found that chromatin organizes along the nuclear periphery, leaving a chromatin-devoid volume at the center of the nucleus, in *Drosophila* myonuclei (Amiad-Pavlov, Lorber et al. 2021). While such studies have primarily characterized chromatin mesoscale organization, how chromatin and the transcription machinery reciprocally shape each other’s spatial distribution in the muscle cells has remained unclear.

In this work, we found that chromatin is organized into clusters, with a dense core which gradually decreases, consistent with other reports (Y. Li et al. 2022; Kant et al. 2024; M. Cremer et al. 2001; Gelléri, Márton et al. 2024; Miron et al. 2020; Nozaki et al. 2017). Peak RNAPII density of each cluster is shifted from the chromatin cluster COM, decreasing both toward the nucleoplasm and toward cluster center, creating a ring-shaped, annular distribution. Whereas this organization is conserved in both control and SUN/*koi* mutants, the size of chromatin clusters increased significantly in the mutants, concomitant with decreased levels of RNAPII associated with each cluster. Global reduction in RNAPII binding to chromatin, as well as increased chromatin repression in SUN/*koi* mutants have previously reported (Amiad Pavlov et al. 2023). However, the high resolution images presented here uniquely reveal the architecture of individual chromatin clusters and their associated RNAPII in wild type and in LINC mutants. This indicates increased size of chromatin clusters accompanied by a significant reduction of RNAPII associated with each chromatin cluster, in the LINC mutants. Furthermore, our findings reveal an asymmetric distribution of chromatin and RNAPII depending on their proximity to the nuclear envelope, and the cluster’s surface orientation. This asymmetry is lost in the SUN/*koi* mutants, highlighting a novel and specific role for the LINC complex in establishing spatial polarity of RNAPII with chromatin, which correlates with the cluster’s distance from the nuclear envelope.

The increase in chromatin clusters size in the LINC mutant (∼ 55% larger clusters, assuming spherical geometry), functionally connects between LINC-dependent binding of chromatin to the nuclear envelope, and its ability to counteract chromatin tendency to self-attract, limiting repressive cluster size (Amiad Pavlov et al. 2023). Previous reports suggested that RNAPII binding to DNA opens chromatin clusters, increasing cluster size (Boettiger et al. 2016; Greiss et al. 2024). Our analysis also indicated an increased size of chromatin clusters associated with RNAPII, that was preserved in both LINC mutants, and BAF knockdown nuclei with. This increase did not lead to elevation of RNAPII levels; instead, it was associated with a decrease in RNAPII levels, possibly implying that the primary event is the enhanced tendency of chromatin to form larger clusters, which then impedes the binding of RNAPII. Enhanced clustering of repressive chromatin and the increased deposition of repressive chromatin marks (e.g. Histone 3K9 and Histone 3K27 methylation), combined with an increase in chromatin binding to repressive transcription factors, including Polycomb and HP1, were both observed in the LINC mutants (Amiad Pavlov et al. 2023) and may explain the enhanced chromatin repression in the LINC mutants and the reduced histone acetylation and decreased binding of RNAPII.

The simulation results provide a mechanistic explanation for the enhanced chromatin clustering, drawing a line between decreased binding of chromatin to the nuclear envelope and an increase in chromatin tendency to self-attract supporting the idea that the binding of chromatin to the nuclear envelope is crucial to maintain the 3D structure of chromatin, and its availability for the transcription machinery. It is noteworthy that the simulations are based on a general assumption that the total amount of chromatin methylation, including LADs and heterochromatin remain unchanged. Furthermore, while the peripheral organization was shown previously to be reduced in the LINC mutant (Amiad Pavlov et al., 2023), it was not reduced quite as much as in the simulation.

The LINC complex interacts with a variety of proteins at the nuclear envelope, including lamins, BAF and LEM domain proteins. Such proteins, including Emerin, and Lamins are involved in chromatin organization and epigenetic modification (Mattout-Drubezki and Gruenbaum 2003; B. Wang, Luo, and Medalia 2025; Marano and Holaska 2025) thus lack of a functional LINC complex possibly affects these interplays. Our previous reports indicated that BAF localization at the nuclear envelope was abolished in LINC mutants (Wang et al. 2018), and here we show that knockdown BAF similarly resulted with the formation of larger chromatin clusters and reduced RNAPII, linking these phenotypes with reduced chromatin binding to the nuclear envelope, independently of the LINC complex. The detection of RNAPII (with an estimated diameter of 150 Å) within the chromatin cluster suggests these structures are sufficiently porous to permit penetration of large protein complexes (Darst, Kubalek, and Kornberg 1989; J. Li et al. 2019). In LINC mutants, the increased cluster size combined with enhanced accumulation of transcriptional repressors may altogether reduce RNAPII accessibility, thereby limiting its penetration into chromatin clusters, as indeed observed. Although we could not accurately quantify the extent of RNAPII reduction in LINC-mutant clusters due to variability in background fluorescence (even within a single nucleus), we observed a significant decrease in RNAPII fluorescence in both the LINC mutants and BAF-KD muscles.

The asymmetry in chromatin and RNAPII distribution observed in clusters proximal to the nuclear envelope likely reflects differences in multiple factors, including density (Gelléri, Márton et al. 2024; Y. Li et al. 2022) transcriptional activity (Peric-hupkes et al. 2010; Robson, Ringel, and Mundlos 2019; Brueckner et al. 2020), the presence of LAD segments (Poleshko et al. 2019), histone marker enrichment (Chen et al. 2018) and altered diffusion coefficients (Daugird et al. 2024). Chromatin adjacent to the nuclear lamina is predominantly composed of repressed facultative heterochromatin typically spanning 200-400 nm in thickness (See et al. 2019; Le Gros et al. 2016;; Li et al. 2022), and beyond this layer chromatin is progressively organized into distinct domains with varying activity patterns. Consistent with our observations, Labade et al. (Labade et al. 2024) reported that RNAPII concentration is low within the first ∼200 nm from the lamina and gradually increases with distance toward the nuclear interior. Since the diameter of chromatin clusters ranges from 100 nm (Miron et al. 2020; Kant et al. 2024; Y. Li et al. 2022) to a few hundred nanometers (Miron et al. 2020; Nozaki et al. 2017; Gelléri et al. 2022), as also observed in this study, it is likely that a given cluster can simultaneously contain both repressed and active regions. Hence, nonuniform chromatin organization and RNAPII can arise on opposite sides of the same cluster. Notably, this asymmetry is lost in LINC mutants, reflecting a link between the proximity of chromatin to the nuclear envelope, and its responsiveness to mechanical inputs, both mediated by the LINC complex. In summary, our findings indicate that LINC-dependent chromatin association with the nuclear envelope prevents chromatin aggregation and transcriptional repression, while promoting asymmetric association of RNAPII with chromatin clusters.

## Supplementary Material

**Supplementary Figure 1:**
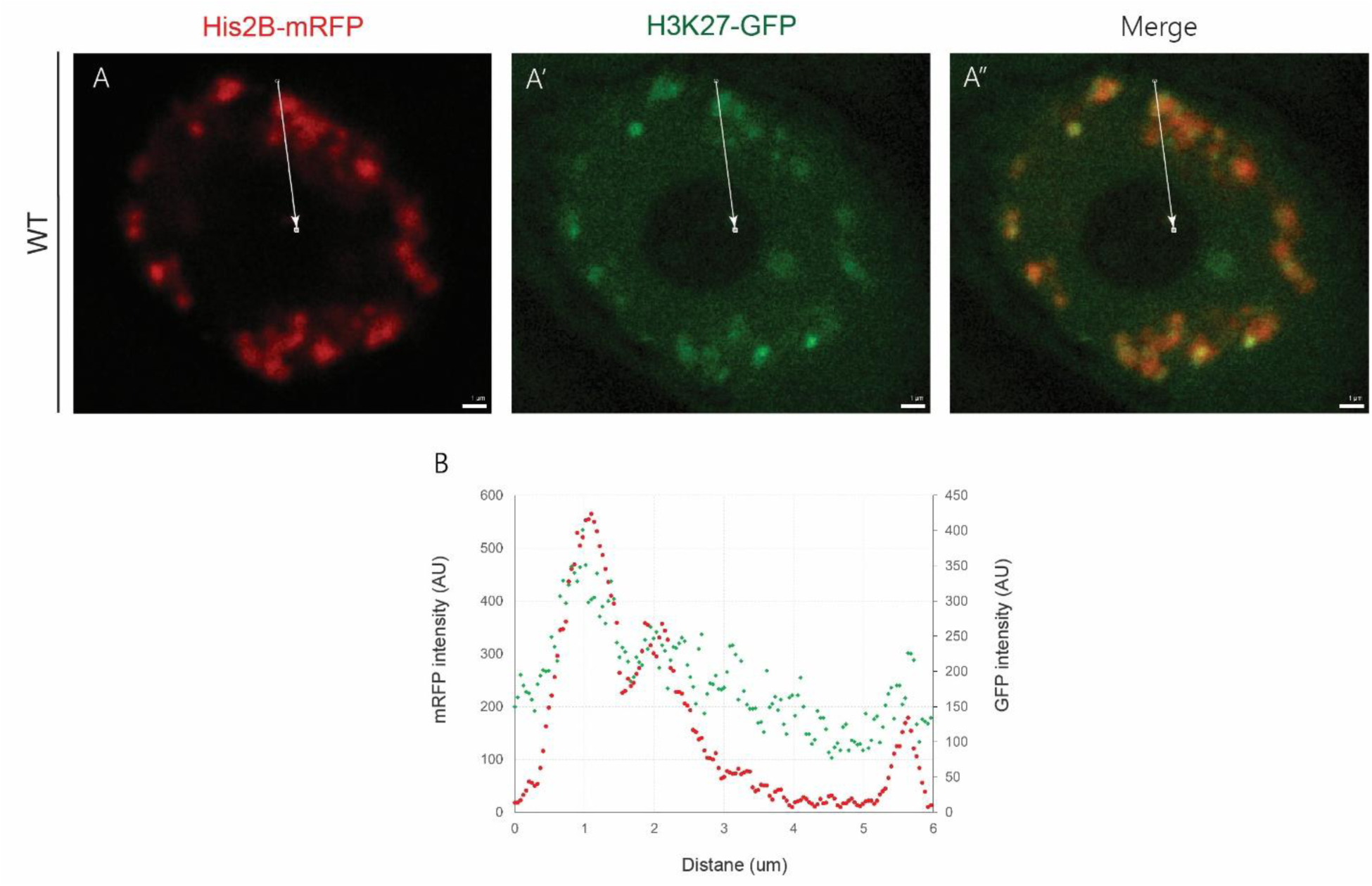
Relative chromatin and H3K27me3 distribution in control muscle nuclei. A-A”: A single optical XY cross sections taken from WT live, intact *Drosophila* larvae. Chromatin (A, red) is labeled with His2B-mRFP, and repressive chromatin H3K27me3 mark is labeled with GFP (A’, green). Their merged image presented (A”). Scale bar=1 μm. B: Intensity profiles of chromatin and H3K27me3 from the nuclear periphery towards the nuclear COM (along the white arrow). H3K27me3 intensity pattern follows chromatin intensity profile, suggesting high repression at the center of the chromatin cluster.

### Computer simulations

Our computer simulations model one chromosome as a multiblock copolymer with three types of repeating blocks: active (type A, euchromatin), inactive (type B, heterochromatin), and LAD (type C, heterochromatin that binds to the lamina). The active block (A) roughly mimics the gene-rich euchromatic regions that contain acetylated nucleosomes. The inactive domains (B) mimic the heterochromatin and type C, lamina-associated, heterochromatin domains. Both the B and C domains are known to be largely gene-inactive since they carry gene-repressed methylated markers on nucleosomes, such as H3k27me3 and H3k9me2/3 and both can be constitutive or facultative (Strom et al. 2017; Larson et al. 2017; Janssen, Colmenares, and Karpen 2018; Sanulli et al. 2019; Zenk et al. 2021) In the spirit of a minimal model, we consider our multiblock copolymer to comprise *N* spherical beads of equal size where *N*_A_, *N*_B_, and *N*_C_ represent the respective number of beads in each block of the active (A), inactive (B), and LAD (C) blocks. The number of A and B blocks is equal, but the number of C blocks can vary (see Supplementary Figure 2).

**Supplementary Figure 2:**
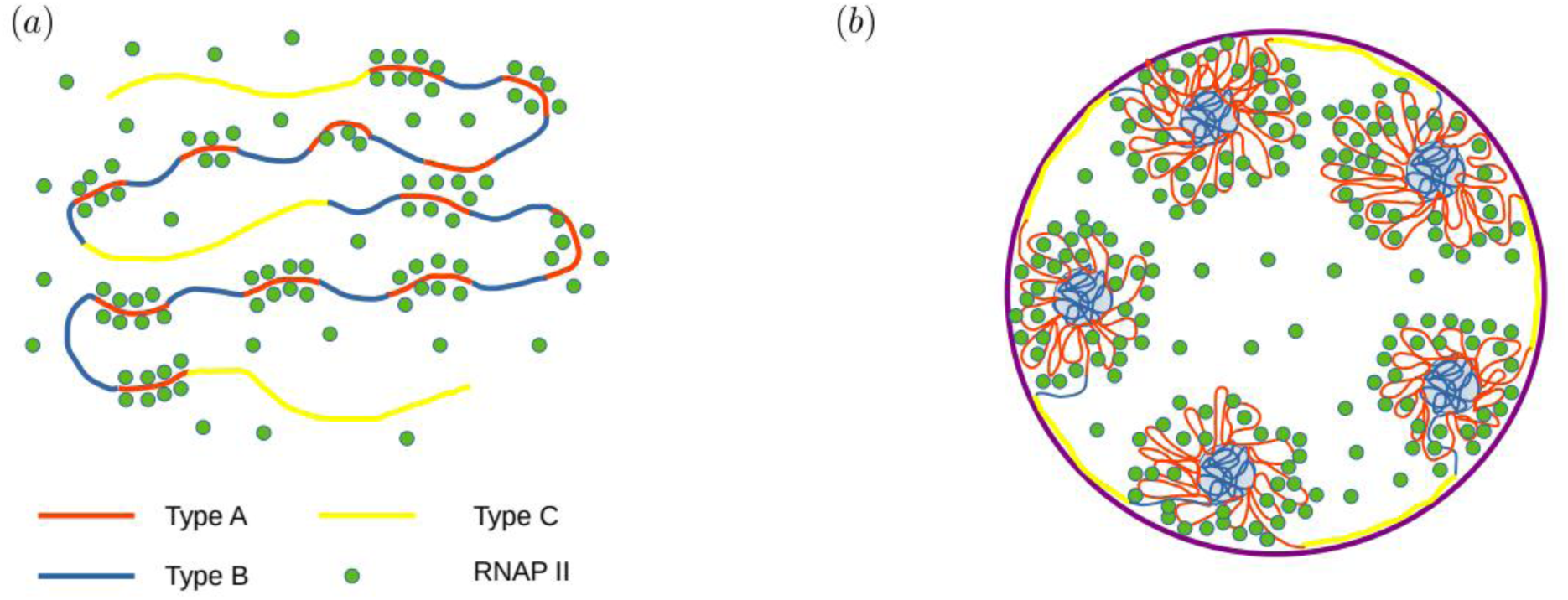
The sketch of model multiblock copolymer model chromatin showing peripheral organization of chromatin micelles. (A) Chromatin is modeled as a multiblock copolymer with periodic block of copolymer domains. Red, blue and yellow domains show type A (euchromatin, EC), type B (heterochromatin, HC) and type C (LAD), respectively. The green beads represent the RNAPII proteins. The periodicity of type A and type B domains is equal, however type C domains have relatively larger periodicity. The RNAPII beads are shown to localize close to the type A domains due to attractive interaction. (B) The spherical confinement mimicking the nuclear lamina is shown with a purple line. LAD domains bind to the confinement wall due to bonded interaction. The differential interactions between the different types of chromatin beads (stronger B-B, relatively weak A-A) lead to miscelle-like organization of chromatin domains consisting of type-B (HC) core and type-A (EC) shell. Such domains are shown to localize in the spatial proximity of the confinement wall due to chain connectivity.

These choices are correlated with (but not identical to) the domains of the in vivo *Drosophila* genome. Thus, one chromosome is modeled by a polymer comprising of *N = n(N_A_ + N_B_) + mN_c_* beads, where *n* is the number of A or B blocks, and *m* is the number of LAD blocks. Motivated by recent *in vivo* and *in vitro* experiments showing phase separation of chromatin from nucleoplasm and buffer (Gibson et al. 2019; Amiad-Pavlov et al. 2021; Amiad Pavlov et al. 2023; T. Cremer et al. 2015), we model chromatin as self-attracting polymer in poor solvent conditions. More detailed *in vivo* studies and microscopy imaging suggest core-shell, micellar organization of chromatin domains. All three types of chromatin phase separate from the nucleoplasm, and within the phase-separated region, the A and B chromatin domains phase separate from each other to form multiple micellar-like clusters. The C-type, LAD chromatin binds to the lamina but since it is topologically connected to the A and B chromatin blocks, these blocks cannot stray too far from the lamina. The individual micelles have a heterochromatin (B) compact core surrounded by a less-condensed shell of euchromatin and RNA polymerase II (RNAP II), due to the differential interaction strengths of AA, BB and AB chromatin contacts. Based on the *in vivo* experiments, all these interactions are significantly stronger than the A, B, or C chromatin interactions with the nucleoplasm which gives rise to the phase separation of all three chromatin types. To replicate the more condensed heterochromatin (B) core, the interaction between inactive monomers is taken to be stronger than active monomers. The interaction hierarchy thus assigns the largest interactions to the chromatin-nucleoplasm and smaller interactions to the AA, BB, and AB contacts. The spatial separation of the A and B chromatin is consistent with previous reports based on chromatin conformation capture approach that established the fact that *in vivo* chromatin organizes into separate domains (Gibson et al. 2019; Amiad-Pavlov et al. 2021; Amiad Pavlov et al. 2023; T. Cremer et al. 2015). In a synthetic context, the selective solvation of copolymer domains, i.e., different A-A and B-B interaction, causes microphase separation of A-type and B-type monomers into various morphologies such as spheres, cylinders, and lamellae. These and other morphologies are due to the competing effect of interfacial tensions and long-range interactions due to chain connectivity. The morphology selected by free energy minimization depends on the interactions, temperature, and the block lengths (Ohta and Kawasaki 1986; Kumar and Safran 2023). A recent study incorporating theoretical modeling and simulations demonstrated that a multiblock copolymer with repeated and contiguous A-type and B-type domains leads to a structure resembling a string of micelles, due to the larger degree of condensation of the B monomers compared to the A, due to stronger BB interactions (Adame-Arana et al. 2023) Applied to the cell nucleus, this allows the less-condensed, active corona of euchromatin (A-type) to be accessible to transcription factors such as RNAPII (Hilbert et al. 2021). It was shown in the previous study that increased binding of RNAPII to the active, A domains, leads to a relative de-condensation of those domains and an increase in the number of denser, inactive (B) cluster cores that are smaller in extent compared to the case of weak binding (Adame-Arana et al. 2023). While this study showed the relative changes in the A and B domains as the binding of RNAPII to the A (euchromatic) domains was varied, it did not include the effects of the C-type LAD domains, their binding to the lamina and how that impacts the organization of the A and B regions.

We thus include A, B, and C types of chromatin as well as spherical beads mimicking RNAPII, which binds to the active, type-A blocks. This attraction to the euchromatin (A) is motivated by the apparent involvement of RNAPII and other transcription factors (Hilbert et al. 2021; Sazer and Schiessel 2018; Brackley and Marenduzzo 2020). Apart from the binding of RNAPII to the A blocks, we do not include their interactions with other transcription factors (“factories”) that associate with active euchromatin or the RNAPII phosphorylation state. The simulations in Fig.6 show the spatial distribution of RNAPII across the entire nucleus as green markers. In addition, the concentration profile of the RNAPII within the chromatin assemblies shows a skewed, bell-shaped distribution with a peak lying in the active euchromatin domain (see Fig. 6 IV). The RNAPII distribution decreases towards the center of the inactive heterochromatin cluster core, eventually approaching a negligible concentration at the center of the core. It is reasonable that the RNAPII distribution is measured to be small in the heterochromatin core of the assemblies, since in addition to the lack of binding to the B blocks, the core formed by those blocks is more condensed with little room for the diffusion of RNAPII. This is why our model only includes binding of the RNAPII beads to the active type-A blocks and short range repulsive (self-avoiding) interactions with the inactive B-blocks. Of course, if we reduced the size of the RNAP II beads they could enter into the B-block, heterochromatin core by diffusion (but not by binding).

The inclusion of the LAD domains (type-C) and their connectivity to the A and B blocks is the new feature of our model compared with the previous study of core(B)-shell(A) micelles of chromatin. The LAD are mostly gene-poor, heterochromatin blocks enriched in repressive histone post translational modifications such as H3K9me2/3 and H3K27me3 (Strom et al. 2017; Larson et al. 2017; Janssen, Colmenares, and Karpen 2018; Sanulli et al. 2019; Zenk et al. 2021). Such domains are known to associate with the nuclear lamina and the inner nuclear membrane (Amiad-Pavlov et al. 2021; Amiad Pavlov et al. 2023) In addition, some of the LAD genomic regions are also known to associate with the nuclear pore complex. We model all the components of the nuclear envelope (lamina, inner nuclear membrane, nuclear pores etc.) by spherical beads that are immobile and result in a spherical confinement of the chromatin and RNAP II; we denote these as confinement beads. The localization of the LAD (C) to the lamina is modeled by a binding interaction with the confinement beads. Since the LAD is topologically connected to the A and B blocks, this leads to peripheral localization of all of the chromatin domains, which additionally have phase separated from the nucleoplasm. Mutations such as *SUN/koi* discussed above, can result in the unbinding of some of the LAD domains from the nuclear envelope. We model this as a reduction in the effective interactions between the C type chromatin and the confinement beads. In the mutants with reduced binding of the C beads to the confinement beads, the LAD fraction that unbinds from the lamina can then merge with inactive (type-B) domains, since both are characterized by mostly methylated nucleosomes. For computational and conceptual simplicity, instead of decreasing the binding strength of the type-C beads to the confinement beads, we coarse-grain the situation in the mutant by reducing the number of C-type beads with a concomitant increase in the number of B-type beads, since both represent heterochromatin.

Brownian dynamics simulation of chromatin peripheral organization and microphase separation *Drosophila* cells: A single chromosome is modeled as a linear polymer with active (A type), inactive (B type), and LAD (C type) monomers. The total number of monomers in the chain, *N = n(N_A_ + N_B_) + mN_C_*, where *n* is the number of inactive and active blocks, *m* is the number of LAD blocks. *N_A_*, *N_B_*, and *N_C_* are the number of active monomers in each block, respectively. In our coarse-grained model, whose focus is the mesoscale organization of the three types of chromatin, each monomer is taken to be a spherical bead of diameter σ representing around 3 kbp of DNA along with 10-12 nucleosomes. We use a bead-spring polymer model where contiguous beads of the polymers are connected by harmonic springs; the *N* beads are thus connected by *N* − 1 harmonic springs. As explained above, the polymer is considered to be self-attracting (poor-solvent conditions) with stronger BB interactions than AA. These non-binding interactions are represented by a truncated Lenard-Jones potential with a cutoff 2.5σ (beyond which the interaction is zero), which has an attractive tail and a repulsive core. The respective strengths of interaction between inactive (BB) and active (AA) beads are taken to be 0.5 and 0.3, respectively. The stronger BB attractions cause those regions to be more condensed, and we took the AA attractions slightly above the critical point of the coil-globule transition of the polymer. The AB attraction between the active and inactive monomers is taken to be the same as the attraction between the AA, active monomers. Such hierarchical interactions lead to a micelle-like core-shell structure observed in the current experiment, as was discussed in the previous simulation study. Unique to our simulation study, we include the LAD (C) monomers that are modeled to have an attractive LJ interaction of strength 0.6 with the confining beads. In addition, we included bonding interactions of the C monomers with the confining beads using a harmonic potential of strength 10*k_B_T/σ^2^* whenever their separation is smaller than 1.5*σ*. Unbinding occurs when the separation of the LAD (C) and confinement beads is larger than 2.5*σ*. This criterion for bonding and un-bonding allows for reversible binding of LAD (C) blocks to the confinement. The LAD (C) bonding to the confining wall results, via the topology of the polymer, an effective localization of the entire polymer and the confinement wall, mimicking nuclear peripheral organization of the chromatin. In addition to chromatin, we include RNAPII modeled as spherical beads that can diffuse in the nucleoplasm but are limited in extent by the spherical confinement. The interaction between these beads is only one of excluded volume, since two beads cannot occupy the same region in the solution, and we do not include RNAPII-RNAPII attractions. Similarly, the interaction between the RNAPII beads and the inactive (B) beads as well as the LAD beads is also only excluded volume. In contrast, we do include an attractive interaction LJ of strength yy between RNAPII beads and active (A) ones. This serves to localize RNAPII near the A beads, which allows for a binding interaction between them. When the separation between the RNAP II and A bead is less than 1.*5σ*, a bond (modeled by a harmonic spring of strength 20*k_B_T/σ^2^*) is formed. The bond is broken once the separation between them increases beyond 2.5*σ*, which allows for equilibrium kinetics. The cumulative effect of both non-bonded and bonded interactions allows for strong localization of the RNAPII beads and the euchromatin (A block) domain. We allow only one RNAPII bead to bind to an A bead (uni-valent binding). When the interactions are large, the RNAPII beads are found right near to active (A) blocks and for large enough binding, coat the entire block. This competes with the AA attractions and for large enough RNAPII binding to the A beads, the self-attraction of the A beads to each other is “screened” (shielded). This results in dilution of the local concentration of the A blocks which even in the absence of RNAPII was less condensed than B blocks. Thus, the A blocks effectively become more soluble in the “poor” nucleoplasm solvent.

**Supplementary Figure 3.**
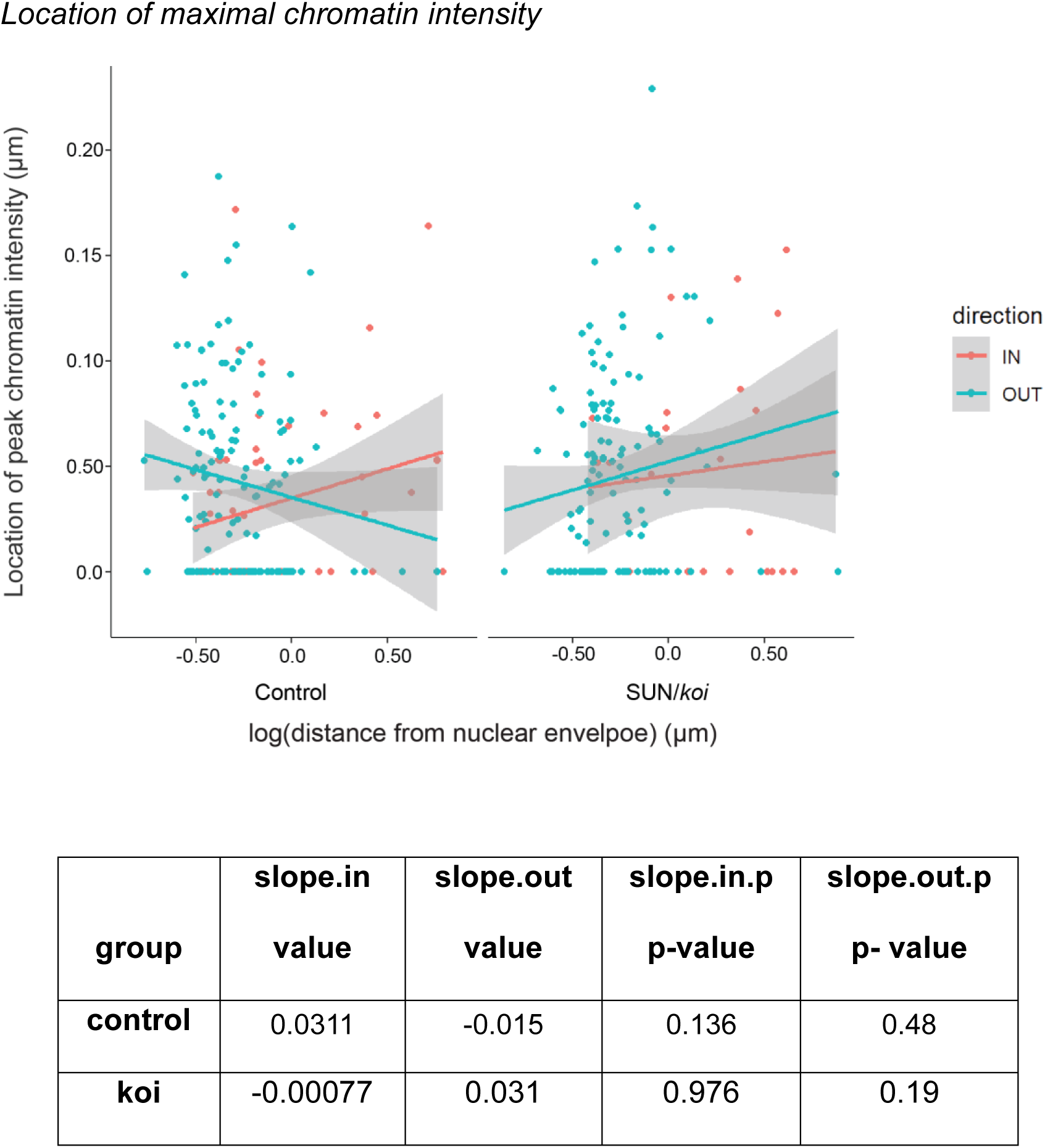
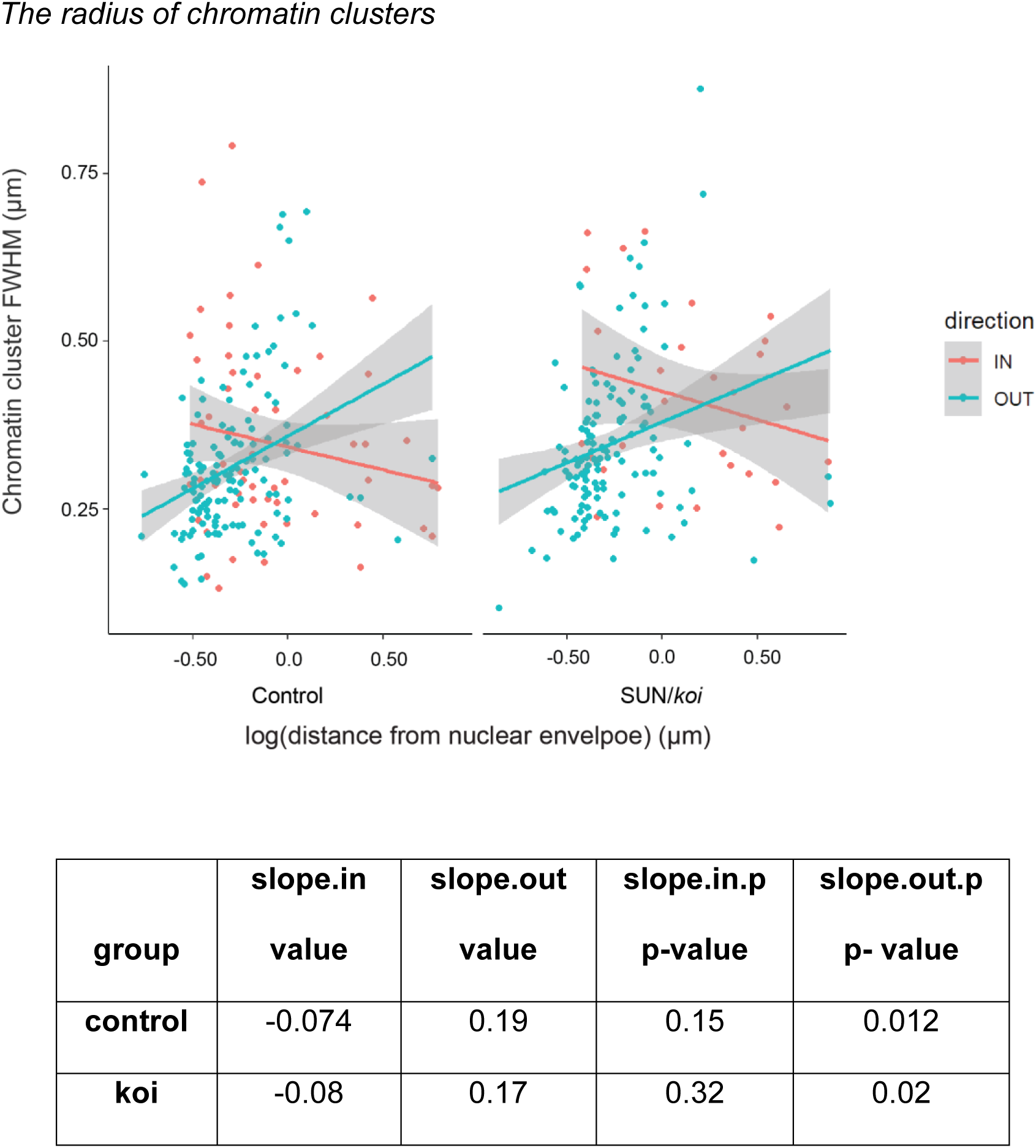
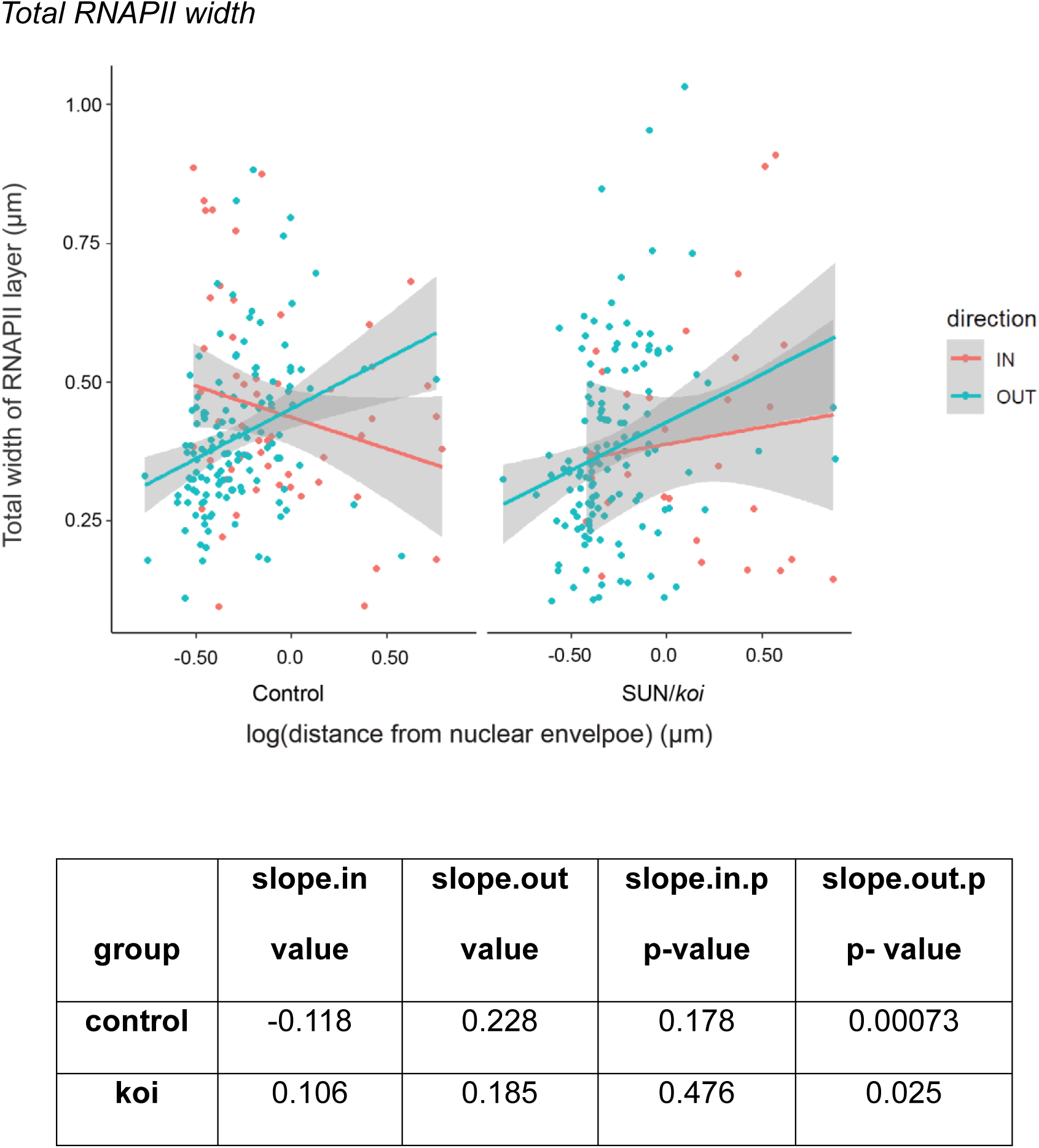
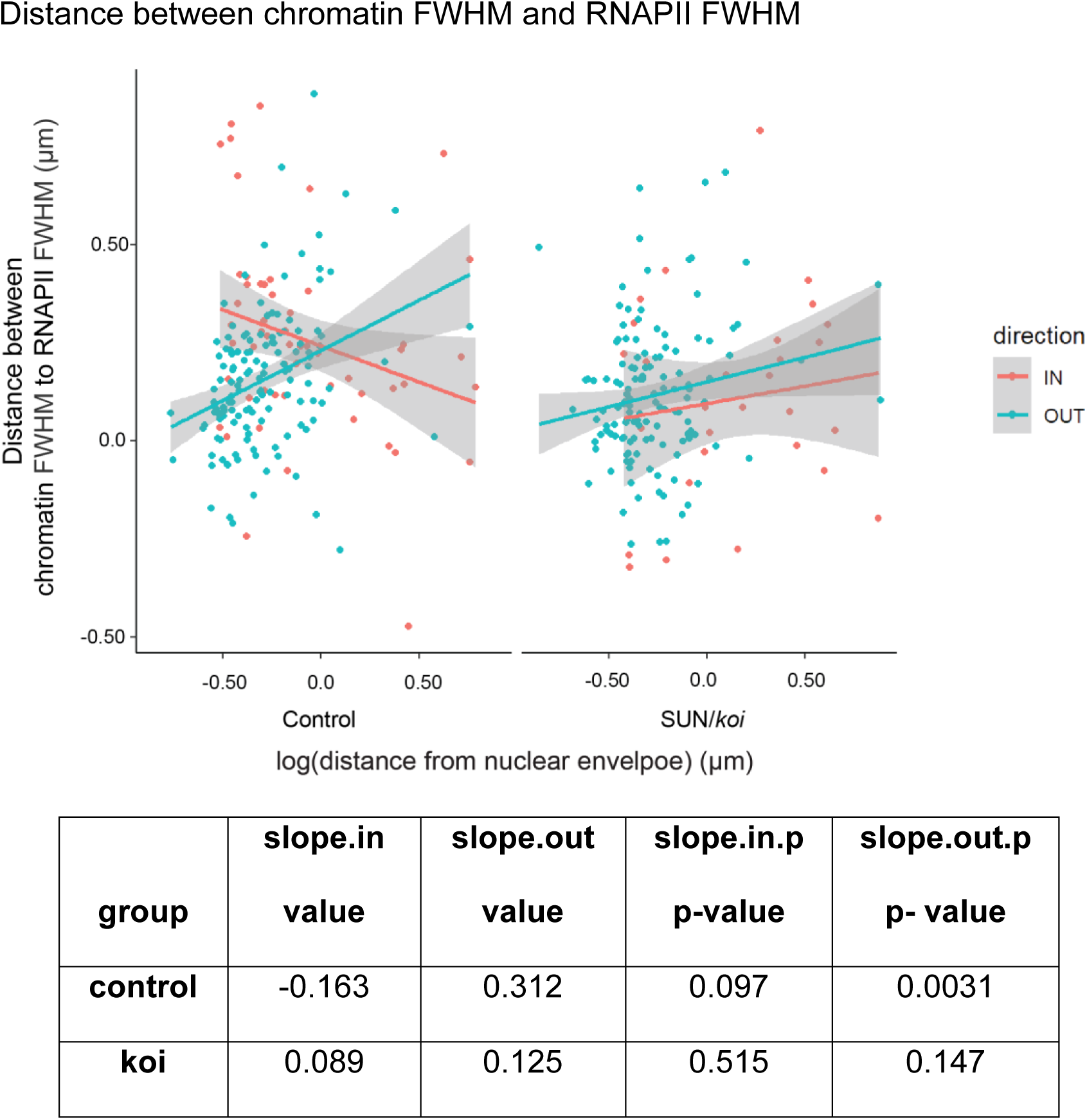
Analyses describing various parameters of the relative chromatin and RNAP II organization per cluster as a function of cluster distance from the nuclear envelope and their surface orientation. For easier readability, the scatterplot trend lines are based on simple linear regression and do not account for the separation to different larvae. Therefore, there are slight differences between the slopes in the plots and the slopes in the mixed effects models.

The analysis revealed that chromatin and RNAPII organization at the surfaces facing the nuclear envelope (denoted OUT) are dependent on the distance from the nuclear envelope, but not at the surface facing the nuclear COM (marked as IN).

## Appendix A: Brownian Dynamics Simulations

The adjacent beads of chromatin polymer are connected by bonds modeled by harmonic springs given by the following potential:

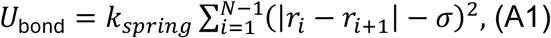

Here, r_i_ and r_i+1_ are the coordinates of the adjacent beads, and bead index varies in the range 1 < i < N − 1, where N is the total number of monomers in the polymer. The equilibrium bond length is equal to σ, which is the bead separation at which the minimum of the harmonic potential occurs. We set the strength of the harmonic potential at a relatively large value, k_spring_ = 100 k_B_T/σ^2^ to ensure that bond fluctuations are not considerable and the bonds are stable. In addition to bonded interaction between the neighboring beads, the pairwise nonbonded interaction between the beads irrespective of their index along the polymer contour is modeled as standard truncated and shifted Lennard-Jones potentials (LJ), U_LJS_(r_ij_) = U_LJ_(r_ij_)+U_LJ_(r_c_), for r_ij_ < r_c_, otherwise U_LJS_(r) = 0. The form of the potential is given by,

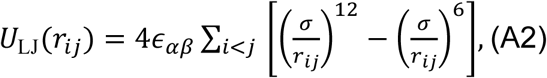

where r_ij_ = |r_i_ − r_j_ | denotes the distance between beads i and j with indices i, j =1, 2, …N. Here, r_i_ and r_j_ are the position vectors of interacting beads, and σ sets the bead diameter. r_c_ represents the cutoff distance at which the LJ potential is truncated and shifted such that the interaction vanishes beyond this separation between the beads. Following the standard practice, we set r_c_ = 2.5σ, which retains both the short-range excluded-volume repulsion and the longer-range attractive tail of the LJ interaction. The potential reaches its minimum at r = 2^1/6^σ, corresponding to the separation at which the attractive and repulsive contributions balance. The strength of interaction potential is given by ɛ_αβ_, where the indices α and β denote the bead type, e.g., active (A), inactive (B) or LAD (C).

Upon increasing the interaction strength from low values, a homopolymer undergoes coil (open conformation) to globule (collapsed conformation) transition in the range 0.2 <ɛαβ < 0.3 k_B_T. To obtain the emergent micelle-like organization of copolymer domains, the interactions between the active, inactive, and LAD monomers are introduced hierarchialy with all interaction strengths chosen above the critical value associated with the coil–globule transition. Specifically, the weakest attractive interaction is assigned between active monomers, ɛ_AA_ = 0.3 k_B_T, corresponding to a marginally collapsed regime. To model the more compact and transcriptionally inactive chromatin domains, the interaction strength between inactive monomers is set to a larger value, ɛ_BB_ = 0.5 k_B_T. The cross-interaction between the active and inactive beads is chosen as ɛ_AB_ = 0.35 k_B_T, ensuring that the active domain remains associated with the inactive domain while forming a sharp interface. In addition, interactions involving lamina-associated monomers are defined such that active–LAD and inactive–LAD interaction strengths are set to ɛ_AC_ = 0.3 k_B_T, and ɛ_BC_ = 0.35 k_B_T, respectively.

The interaction between the chromatin polymer and the confining boundary is primarily mediated through the lamina-associated (LAD) blocks along the polymer chain. These interactions are modeled using a truncated and shifted Lennard–Jones (LJ) potential, analogous to the bead–bead interaction described in Eq. A2. The wall–bead interaction potential is defined as: U_wall_(r_ij_) = ULJwall(rij) + U_LJwall_(r_c_), for r_ij_ < r_c_, otherwise U_wall_(r_ij_) = 0, where r_c_ is the cutoff distance. The unshifted LJ interaction between polymer beads and wall beads is given by,

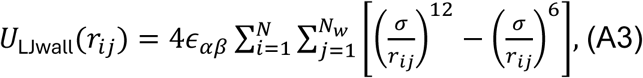

with r_ij_ =| r_i_ − r_wj_| denoting the separation between polymer bead i and wall bead j.

Here, r_i_ and r_wj_ represent the position vectors of the polymer beads and the confinement wall beads, respectively, with indices i = 1, 2, …,N, and j = 1, 2, 3, …,N_w_. Please note that the confinement wall is an assembly of a total N_w_ spherical beads of the same size as monomers of the chain. The cutoff distance for the wall–bead LJ interaction is chosen as r_c_ = 2.5σ. retaining both the repulsive core and attractive tail of the interaction. The strength of attraction between lamina-associated (LAD) monomers and the confinement wall is set to ɛ_CW_ = 0.6 k_B_T, corresponding to strong attraction with the nuclear periphery. Apart from LAD monomers, interactions between active and inactive monomers with the confinement wall are modeled using a purely repulsive Weeks–Chandler–Andersen (WCA) potential. This interaction is commonly employed to enforce excluded-volume (self-avoiding) effects between particles. The WCA potential corresponds to the repulsive part of the Lennard–Jones interaction given as Eq. A3, and obtained by truncating and shifting the LJ potential at its minimum, r_c_^w^ = 2^1/6^σ, ensuring continuity of both the potential and force at the cutoff. Thus, the potential becomes: U_WCAwall_(r_ij_) = U_LJwall_(r_ij_) + U_LJwall_(r_c_^w^), for r_ij_ < r_c_^w^, otherwise U_wall_(r_ij_) = 0. The notations have the same meaning as given above. The interaction strength is set to ɛ_AW_ = ɛ_BW_ = 1 k_B_T, providing a purely steric repulsion between active or inactive monomers and the confining boundary.

In addition to the nonbonded interaction between LAD monomers and the confinement wall, we introduce the bonded interaction between them. Each LAD monomer is allowed to bind to at most one bead of the confinement wall. A bond is formed when the spatial separation between the LAD monomer and confinement wall is less than a distance, r_bond_ = 1.5σ. For strong binding, the strength of the harmonic potential is fixed at K = 10 k_B_T/σ^2^. After formation of the bond, the bond length adopts an equilibrium value r_0_ = σ around which small fluctuations occur. In addition, the bond dissociation is also permitted when the separation between the LAD monomers and the wall exceed beyond the cutoff value, r_break_. We fix this distance to be same as the LJ cutoff r_c_ at which the LJ potential vanishes; thus r_break_ = 2.5σ. The formation and dissociation of bonds are implemented using the built-in Monte Carlo (MC) bonding algorithm in the LAMMPS simulation package (Plimpton 1995; Thompson et al. 2022) which is described by

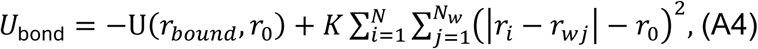

Here, r_i_ and r^w^_j_ are the coordinates of LAD monomers and confinement wall beads, respectively. For their separation less than the chosen cutoff, i.e., |r_i_ − r^w^_j_| < r_break_, the above potential is applicable, otherwise it vanishes. The quantity U_0(rbond, r0)_ denotes the bond formation energy.

In addition to the chromatin polymer, we introduce transcription factor RNA polymerase II (RNAPII) beads within the spherical confinement. These RNAPII complexes are modeled as spherical particles of diameter σ, identical to the monomers comprising the chromatin polymer. RNAPII beads are assumed to preferentially associate with transcriptionally active (euchromatic) regions of chromatin. The interactions between RNAP II beads and inactive monomers, as well as between RNAP II beads and lamina-associated (LAD) monomers, are modeled using a purely repulsive Weeks–Chandler–Andersen (WCA) potential, identical in form to that defined in Eq. A3. In this case, the only modification to Eq. A3 is that the summation index j runs over RNAPII beads, j = 1, 2, 3…,Np, where Np denotes the total number of RNAPII particles in the system. In contrast, the interaction between RNAPII beads and active chromatin monomers is taken to be attractive. This interaction is modeled using the standard Lennard–Jones potential defined in Eq. A2. The strength of the RNAP II–active interaction is fixed at an intermediate value, ɛ_AP_ = 0.4, which promotes localization of RNAPII beads in the vicinity of active chromatin regions. This choice facilitates frequent encounters between RNAPII beads and active monomers, thereby enhancing the probability of subsequent binding events.

Further, in addition to the nonbonded Lennard–Jones interactions, we incorporate reversible bonded interactions between RNAPII beads and chromatin monomers. In contrast to the LAD–wall case, the valency of the RNAP II –chromatin bonded interaction is fixed at one, allowing each RNAP II bead to form bonds with only one chromatin monomer and *vice versa*. Bond formation between a RNAP II bead and a chromatin monomer occurs in a manner analogous to the LAD–wall bonded interaction described in Eq. A4. Specifically, when the separation between a RNAP II bead and an active chromatin monomer falls below a capture distance *r*_bond_ = 1.5*σ*, a harmonic bond is created. The bonded inter-action is described by a harmonic potential with a spring constant, *K* = 20 *k*_B_*T/a*^2^, leading to a strong bond allowing small fluctuations around equilibrium. Bond dissociation is per-mitted when the separation between the RNAP II bead and the chromatin monomer exceeds a breaking distance *r*_break_. As in the LAD–wall case, we set *r*_break_ = *r_c_* = 2.5*σ*, ensuring consistency between bonded and nonbonded interaction length scales. Apart from the differences in bead type and particle number, all other aspects of the bond formation and dis-sociation protocol are identical to those employed for LAD–wall bonding. The total number of RNAP II beads is fixed at *N_p_* = 10000. This choice ensures that most active chromatin monomers are accompanied by at least one RNAP II bead, while simultaneously avoiding excessive crowding within the confined volume. Interactions between RNAP II beads and the confining wall are modeled using a purely repulsive Weeks–Chandler–Andersen (WCA) potential, identical to that used for active monomer–wall interactions. We incorporate all the interactions (bonded and nonbonded) involving RNAP II beads as *U*_pol_. The total potential energy of the chromatin-RNAP II system is given by

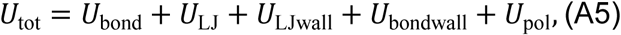

As the entire system is confined within a spherical boundary, the radius of confinement *R_c_* is determined by fixing the global volume fraction *ϕ* of all the beads. We choose *ϕ* = 0.05, ensuring that the system corresponds to a dilute regime in which beads diffuse freely without significant steric crowding:

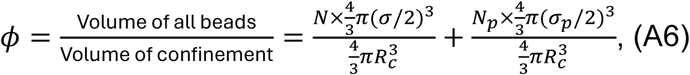

where *N* and *N_p_* are the total number of beads in the chromatin chain, and RNAP II beads, respectively. In our current simulations, we fix *N* = 50, 000 and *N_p_* = 10000. The diameters of chromatin beads and RNAP II beads are given as *σ* and *σ_p_*, respectively. We set *σ_m_* = *σ_p_* = 1 such that all the beads have the same size. Using the global volume fraction *ϕ* = 0.05, and following Eq. (A6), we obtain *R_c_* = 51.36. This choice of volume fraction ensures that the confined system remains dilute, with beads occupying only a small fraction of the available nuclear volume, thereby minimizing crowding effects.

We perform the molecular dynamics simulations using the standard velocity-Verlet algorithm implemented by LAMMPS package. This method solves Newton’s equation of motion for the beads embedded in a solvent, providing thermal noise and dissipation at equilibrium. All the computations are conducted in the overdamped limit. The time step for the simulations is adopted to be *δt* = 0.01*τ*, where 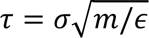 is the characteristic time. The mass of the beads is fixed at unity. The thermostat temperature is kept constant at *T* = *ɛ/k_B_*, using the Langevin thermostat incurring a damping constant 10*τ*. Here *ɛ* and *k_B_* are the strength of the LJ potential and the Boltzmann constant, respectively. For simplicity, we set *ɛ* = 1, and *k_B_* = 1. In this scheme, pure diffusion of a bead takes a time *γσ*^2^*/k_B_T* to diffuse over its diameter *σ*, which is equal to the characteristic time. The simulation runs over more than 10^7^*τ*, ensuring the configurations are equilibrated. The presented data are captured over equilibrated configurations.

## Acknowledgments

We thank K. Hiroshi (Osaka University, Japan), K. Furukawa (Niigata University, Japan), Patrick O’Farrell (UCSF, USA), and the Bloomington Stock Center for providing fly lines. We thank the devoted imaging team, Yoseph Addadi, Tatiana Smirnova, and Inna Goliand from the Advanced Optical Imaging Unit, de Picciotto-Lesser Cell Observatory unit, at the Moross Integrated Cancer Center Life Sciences Core Facilities, Weizmann Institute of Science. We acknowledge Samuel Gelman from the Bioinformatics unit at the Life Sciences Core Facilities for the help with the initial numerical analysis. We are grateful to Gaurav Bajpai for many useful discussions about the simulation technique.

## Author contributions

D.L. and T.V. conceptualized the experiments. A.K. and S.S. conceptualized the theoretical model. D.L. performed the experiments and image acquisition. I.A. conceptualized the algorithmic pipeline and wrote and applied the Python code. I.A and D.L. performed the image analysis, and R.R performed the statistical analysis. A.K. and S.S. performed model computer simulations and analysis. S.S. reviewed and edited the manuscripts. T.V. and D.L. wrote the manuscript.

## Data and materials availability

All data needed to evaluate the conclusions in the paper are present in the paper and/or the Supplementary Materials. Additional data related to this paper may be requested from the authors.

## References

Adame-Arana, Omar, Gaurav Bajpai, Dana Lorber, Talila Volk, and Samuel Safran. 2023. “Regulation of Chromatin Microphase Separation by Binding of Protein Complexes.” ELife 12: 1–21. 10.7554/eLife.82983.

Almassalha, Luay M., Marcelo Carignano, Emily Pujadas Liwag, Wing Shun Li, Ruyi Gong, Nicolas Acosta, Cody L. Dunton, et al. 2025. “Chromatin Conformation, Gene Transcription, and Nucleosome Remodeling as an Emergent System.” Science Advances 11: eadq6652. 10.1126/sciadv.adq6652.

Amiad-Pavlov, Daria, Dana Lorber, Gaurav Bajpai, Adriana Reuveny, Francesco Roncato, Ronen Alon, Samuel Safran, and Talila Volk. 2021. “Live Imaging of Chromatin Distribution in Muscle Nuclei Reveals Novel Principles of Nuclear Architecture and Chromatin Compartmentalization.” Science Advances 7 (23).

Amiad Pavlov, Daria, C. P. Unnikannan, Dana Lorber, Gaurav Bajpai, Tsviya Olender, Elizabeth Stoops, Adriana Reuveny, Samuel Safran, and Talila Volk. 2023. “The LINC Complex Inhibits Excessive Chromatin Repression.” Cells 12 (6): 932. 10.3390/cells12060932.

Andrés, Vicente, and José M. González. 2009. “Role of A-Type Lamins in Signaling, Transcription, and Chromatin Organization.” Journal of Cell Biology 187 (7): 945–57. 10.1083/jcb.200904124.

Auld, A. L., and E. S. Folker. 2016. “Nucleus-Dependent Sarcomere Assembly Is Mediated by the LINC Complex.” Molecular Biology of the Cell 27: 2351–59. 10.1091/mbc.E16-01-0021.

Bajpai, Gaurav, Daria Amiad Pavlov, Dana Lorber, Talila Volk, and Samuel Safran. 2021. “Mesoscale Phase Separation of Chromatin in the Nucleus.” ELife 10: 1–27. 10.7554/elife.63976.

Banigan, Edward J., Wen Tang, Aafke A. van den Berg, Roman R. Stocsits, Gordana Wutz, Hugo B. Brandão, Georg A. Busslinger, Jan-Michael Peters, and Leonid A. Mirny. 2023. “Transcription Shapes 3D Chromatin Organization by Interacting with Loop Extrusion.” PNAS 120 (11). 10.1073/pnas.

Barton, Lacy J., Alexey A. Soshnev, and Pamela K. Geyer. 2015. “Networking in the Nucleus: A Spotlight on LEM-Domain Proteins.” Current Opinion in Cell Biology 34: 1–8. 10.1016/j.ceb.2015.03.005.

Bemmel, Joke G. van, Ludo Pagie, Ulrich Braunschweig, Wim Brugman, Wouter Meuleman, Ron M. Kerkhoven, and Bas van Steensel. 2010. “The Insulator Protein SU(HW) Fine-Tunes Nuclear Lamina Interactions of the Drosophila Genome.” PLoS ONE 5 (11). 10.1371/journal.pone.0015013.

Bersaglieri, Cristiana, and Raffaella Santoro. 2023. “Methods for Mapping 3D-Chromosome Architecture around Nucleoli.” Current Opinion in Cell Biology 81: 102171. 10.1016/j.ceb.2023.102171.

Boettiger, Alistair N., Bogdan Bintu, Jeffrey R. Moffitt, Siyuan Wang, Brian J. Beliveau, Geoffrey Fudenberg, Maxim Imakaev, Leonid A. Mirny, Chao Ting Wu, and Xiaowei Zhuang. 2016. “Super-Resolution Imaging Reveals Distinct Chromatin Folding for Different Epigenetic States.” Nature 529: 418–22. 10.1038/nature16496.

Bouzid, Tasneem, Eunju Kim, brandon h. Riehl, Amir Monemian Esfahani, Jordan Rosenbohm, Ruiguo Yang, Bin Duan, and Jung yul Lim. 2019. “The LINC Complex, Mechanotransduction,And.” Journal of Biological Engineering 13 (68).

Brackley, C A, and D Marenduzzo. 2020. “Bridging-Induced Microphase Separation: Photobleaching Experiments, Chromatin Domains and the Need for Active Reactions.” Briefings in Functional Genomics 19 (2): 111–18. 10.1093/bfgp/elz032.

Bradski, Gary. 2000. “The OpenCV Library.” Dr. Dobb’s Journal: Software Tools for the Professional Programme 25 (11): 120–23.

Brueckner, Laura, Peiyao A Zhao, Tom van Schaik, Christ Leemans, Jiao Sima, Daniel Peric-Hupkes, David M Gilbert, and Bas van Steensel. 2020. “Local Rewiring of Genome–Nuclear Lamina Interactions by Transcription.” The EMBO Journal 39 (6): 1–17. 10.15252/embj.2019103159.

Cartwright, Sarah, and Iakowos Karakesisoglou. 2014. “Nesprins in Health and Disease.” Seminars in Cell and Developmental Biology 29: 169–79. 10.1016/j.semcdb.2013.12.010.

Chen, Yu, Yang Zhang, Yuchuan Wang, Liguo Zhang, Eva K. Brinkman, Stephen A. Adam, Robert Goldman, Bas Van Steensel, Jian Ma, and Andrew S. Belmont. 2018. “Mapping 3D Genome Organization Relative to Nuclear Compartments Using TSA-Seq as a Cytological Ruler.” Journal of Cell Biology 217 (11): 4025–48. 10.1083/jcb.201807108.

Cho, Chun-Yi, James P. Jr Kemp, Robert J. Duronio, and Patrick H. O’Farrell. 2022. “Coordinating Transcription and Replication to Mitigate Their Conflicts in Early Drosophila Embryos.” Cell Reports 41 (3): 1–24. 10.1016/j.celrep.2022.111507.Coordinating.

Cho, Won-ki, Namrata Jayanth, Brian P English, Takuma Inoue, J Owen Andrews, William Conway, Jonathan B Grimm, et al. 2016. “RNA Polymerase II Cluster Dynamics Predict MRNA Output in Living Cells.” ELife 5: 1–31. 10.7554/eLife.13617.

Cho, Won Ki, Jan Hendrik Spille, Micca Hecht, Choongman Lee, Charles Li, Valentin Grube, and Ibrahim I. Cisse. 2018. “Mediator and RNA Polymerase II Clusters Associate in Transcription-Dependent Condensates.” Science 361 (6400): 412–15. 10.1126/science.aar4199.

Chowdhary, Surabhi, Sarah Paracha, Lucas Dyer, and David Pincus. 2025. “Emergent 3D Genome Reorganization from the Stepwise Assembly of Transcriptional Condensates.” BioRxiv.

Coninck, Dennis de, Thomas H. Schmidt, Jan Gero Schloetel, and Thorsten Lang. 2018. “Packing Density of the Amyloid Precursor Protein in the Cell Membrane.” Biophysical Journal 114: 1128–41. 10.1016/j.bpj.2018.01.009.

Cremer, M., J. V. Hase, T. Volm, A. Brero, G. Kreth, J. Walter, C. Fischer, I. Solovei, C. Cremer, and T. Cremer. 2001. “Non-Random Radial Higher-Order Chromatin Arrangements in Nuclei of Diploid Human Cells.” Chromosome Research 9 (7): 541–67. 10.1023/A:1012495201697.

Cremer, Marion, Volker J. Schmid, Felix Kraus, Yolanda Markaki, Ines Hellmann, Andreas Maiser, Heinrich Leonhardt, Sam John, John Stamatoyannopoulos, and Thomas Cremer. 2017. “Initial High-Resolution Microscopic Mapping of Active and Inactive Regulatory Sequences Proves Non-Random 3D Arrangements in Chromatin Domain Clusters.” Epigenetics and Chromatin 10: 1–17. 10.1186/s13072-017-0146-0.

Cremer, Thomas, and Christoph Cremer. 2001. “Chromosome Territories, Nuclear Architecture and Gene Regulation in Mammalian Cells.” Nature Reviews Genetics 2 (April): 292–301. 10.1038/35066075.

Cremer, Thomas, Marion Cremer, Barbara Hübner, Hilmar Strickfaden, Daniel Smeets, Jens Popken, Michael Sterr, Yolanda Markaki, Karsten Rippe, and Christoph Cremer. 2015. “The 4D Nucleome: Evidence for a Dynamic Nuclear Landscape Based on Co-Aligned Active and Inactive Nuclear Compartments.” FEBS Letters 589 (20): 2931–43. 10.1016/J.FEBSLET.2015.05.037.

Darst, Seth A., Elizabeth W. Kubalek, and Roger D. Kornberg. 1989. “Three-Dimensional Structure of Escherichia Coli RNA Polymerase Holoenzyme Determined by Electron Crystallography.” Nature 340: 730–32. 10.1038/340730a0.

Daugird, Timothy A, Yu Shi, Katie L Holland, Hosein Rostamian, Zhe Liu, Luke D Lavis, Joseph Rodriguez, Brian D Strahl, and Wesley R Legant. 2024. “Correlative Single Molecule Lattice Light Sheet Imaging Reveals the Dynamic Relationship between Nucleosomes and the Local Chromatin Environment.” Nature Communications 15: 1–20. 10.1038/s41467-024-48562-0.

Dekker, Job, Betul Akgol Oksuz, Yang Zhang, and Ye Wang. 2024. “An Integrated View of the Structure and Function of the Human 4D Nucleome.” BioRxiv, 1–67.

Dubochet, J, and N Sartori Blanc. 2001. “The Cell in Absence of Aggregation Artifacts.” Micron 32: 91–99.

Dupont, Claire, Dhanvantri Chahar, Antonio Trullo, Thierry Gostan, Caroline Surcis, Charlotte Grimaud, Daniel Fisher, Robert Feil, and David Llères. 2023. “Evidence for Low Nanocompaction of Heterochromatin in Living Embryonic Stem Cells.” The EMBO Journal 42 (12). 10.15252/embj.2021110286.

Elhanany-Tamir, Hadas, Yanxun V. Yu, Miri Shnayder, Ankit Jain, Michael Welte, and Talila Volk. 2012. “Organelle Positioning in Muscles Requires Cooperation between Two KASH Proteins and Microtubules.” Journal of Cell Biology 198 (5): 833–46. 10.1083/jcb.201204102.

Elimelech, Aviv, and Ramon Y. Birnbaum. 2020. “From 3D Organization of the Genome to Gene Expression.” Current Opinion in Systems Biology 22: 22–31. 10.1016/j.coisb.2020.07.006.

Fursova, Nadezda A., and Daniel R. Larson. 2024. “Transcriptional Machinery as an Architect of Genome Structure.” Current Opinion in Structural Biology 89. 10.1016/j.sbi.2024.102920.

Gelléri, Márton, Michael Sterr, Hilmar Strickfaden, Christoph Cremer, Thomas Cremer, and Marion Cremer. 2024. “Space-Time Dynamics of Genome Replication Studied with Super-Resolved Microscopy.” BioRxiv, 1–26.

Gelléri, Márton, Shih-Ya Chen, Aleksander Szczurek, Barbara Hübner, Michael Sterr, Jan Neumann, Ole Kröger, et al. 2022. “True-to-Scale DNA-Density Maps Correlate With Major Accessibility Differences Between Active and Inactive Chromatin.” SSRN Electronic Journal. 10.2139/ssrn.4162083.

Gelléri, Márton, Shih Ya Chen, Barbara Hübner, Jan Neumann, Ole Kröger, Filip Sadlo, Jorg Imhoff, et al. 2023. “True-to-Scale DNA-Density Maps Correlate with Major Accessibility Differences between Active and Inactive Chromatin.” Cell Reports 42 (112567). 10.1016/j.celrep.2023.112567.

Georgiev, Tihomir, Mikhail Svirin, Enrique Jaimovich, and Rainer H.A. Fink. 2015. “Localized Nuclear and Perinuclear Ca2+signals in Intact Mouse Skeletal Muscle Fibers.” Frontiers in Physiology 6. 10.1088/0031-9155/61/2/758.

Gibson, Bryan A., Lynda K. Doolittle, Maximillian W.G. Schneider, Liv E. Jensen, Nathan Gamarra, Lisa Henry, Daniel W. Gerlich, Sy Redding, and Michael K. Rosen. 2019. “Organization of Chromatin by Intrinsic and Regulated Phase Separation.” Cell 179 (2): 470–484.e21. 10.1016/j.cell.2019.08.037.

Gilchrist, Daniel A, Gilberto Dos Santos, David C Fargo, Bin Xie, Yuan Gao, Leping Li, and Karen Adelman. 2010. “Pausing of RNA Polymerase II Disrupts DNA-Specified Nucleosome Organization to Enable Precise Gene Regulation.” Cell 143: 540–51. 10.1016/j.cell.2010.10.004.

Goronzy, Isabel N., Sofia A. Quinodoz, Joanna W. Jachowicz, Noah Ollikainen, Prashant Bhat, and Mitchell Guttman. 2022. “Simultaneous Mapping of 3D Structure and Nascent RNAs Argues against Nuclear Compartments That Preclude Transcription.” Cell Reports 41. 10.1016/j.celrep.2022.111730.

Greiss, Ferdinand, Shirley S Daube, Vincent Noireaux, and Roy Bar-ziv. 2024. “Transcription-Dependent Swelling of a Transplanted Chromosome in an Artificial Cell.” BioRxiv, 1–25.

Gros, Mark A. Le, E. Josephine Clowney, Angeliki Magklara, Angela Yen, Eirene Markenscoff-Papadimitriou, Bradley Colquitt, Markko Myllys, Manolis Kellis, Stavros Lomvardas, and Carolyn A. Larabell. 2016. “Soft X-Ray Tomography Reveals Gradual Chromatin Compaction and Reorganization during Neurogenesis In Vivo.” Cell Reports 17: 2125–36. 10.1016/j.celrep.2016.10.060.

Gruenbaum, Yosef, and Roland Foisner. 2015. “Lamins: Nuclear Intermediate Filament Proteins with Fundamental Functions in Nuclear Mechanics and Genome Regulation.” Annual Review of Biochemistry 84 (1): 131–64. 10.1146/annurev-biochem-060614-034115.

Guelen, Lars, Ludo Pagie, Emilie Brasset, Wouter Meuleman, Marius B. Faza, Wendy Talhout, Bert H. Eussen, et al. 2008. “Domain Organization of Human Chromosomes Revealed by Mapping of Nuclear Lamina Interactions.” Nature 453: 948–51. 10.1038/nature06947.

Haaf, Thomas, and David C Ward. 1996. “Inhibition of RNA Polymerase II Transcription Causes Chromatin Decondensation, Loss of Nucleolar Structure, and Dispersion of Chromosomal Domains.” Experimental Cell Research 224: 163–73.

Harris, Charles R, K Jarrod Millman, Stéfan J Van Der Walt, Ralf Gommers, Pauli Virtanen, David Cournapeau, Eric Wieser, et al. 2020. “Array Programming with NumPy.” Nature 585 (September): 357–62. 10.1038/s41586-020-2649-2.

Haws, Spencer A., Zoltan Simandi, R. Jordan Barnett, and Jennifer E. Phillips-Cremins. 2022. “3D Genome, on Repeat: Higher-Order Folding Principles of the Heterochromatinized Repetitive Genome.” Cell 185: 2690–2707. 10.1016/j.cell.2022.06.052.

Hilbert, Lennart, Yuko Sato, Ksenia Kuznetsova, Tommaso Bianucci, Hiroshi Kimura, Frank Jülicher, Alf Honigmann, Vasily Zaburdaev, and Nadine L. Vastenhouw. 2021. “Transcription Organizes Euchromatin via Microphase Separation.” Nature Communications 12 (1): 1360. 10.1038/s41467-021-21589-3.

Hunter, John D. 2007. “Matplotlib: A 2D Graphics Environment.” Computing in Science & Engineering, 90–95.

Imada, Takashi, Takeshi Shimi, Ai Kaiho, Yasushi Saeki, and Hiroshi Kimura. 2021. “RNA Polymerase II Condensate Formation and Association with Cajal and Histone Locus Bodies in Living Human Cells.” Genes to Cells 26: 298–312. 10.1111/gtc.12840.

Irgen-Gioro, Shawn, Shawn Yoshida, Victoria Walling, and Shasha Chong. 2022. “Fixation Can Change the Appearance of Phase Separation in Living Cells.” ELife 11: 1–24.

Jahed, Zeinab, and Mohammad RK Mofrad. 2019. “The Nucleus Feels the Force, LINCed in or Not !” Current Opinion in Cell Biology 58: 114–19.

Janssen, Aniek, Serafin U Colmenares, and Gary H Karpen. 2018. “Heterochromatin : Guardian of the Genome.” Annual Review OfCell and Developmental Biology 34: 265–88.

Javed, Khayam, Jerome Jullien, Gaurav Agarwal, Nicola Lawrence, Richard Butler, Pantelis Savvas Ioannou, Farhat Nazir, and J. B. Gurdon. 2022. “DNA-Induced Spatial Entrapment of General Transcription Machinery Can Stabilize Gene Expression in a Nondividing Cell.” Proceedings of the National Academy of Sciences of the United States of America 119. 10.1073/pnas.2116091119.

Kant, Aayush, Zixian Guo, Vinayak Vinayak, Maria Victoria Neguembor, Wing Shun Li, Vasundhara Agrawal, Emily Pujadas, et al. 2024. “Active Transcription and Epigenetic Reactions Synergistically Regulate Meso-Scale Genomic Organization.” Nature Communications 15: 4338. 10.1038/s41467-024-48698-z.

King, Megan C. 2023. “Dynamic Regulation of LINC Complex Composition and Function across Tissues and Contexts” 597: 2823–32. 10.1002/1873-3468.14757.

Kracklauer, Martin P., Susan M.L. Banks, Xuanhua Xie, Yaning Wu, and Janice A. Fischer. 2007. “Drosophila Klaroid Encodes a SUN Domain Protein Required for Klarsicht Localization to the Nuclear Envelope and Nuclear Migration in the Eye.” Fly 1 (2): 75–85. 10.4161/fly.4254.

Kumar, Amit, and Samuel A Safran. 2023. “Fluctuations and Shape Dependence of Microphase Separation in Systems with Long-Range Interactions.” Physical Review Letters 131 (258401): 1–7. 10.1103/PhysRevLett.131.258401.

Labade, Ajay S., Caroline Comenho Zachary D. Chiang, Paul L. Reginato, Andrew C. Payne, Andrew S. Earl, Rojesh Shrestha, Fabiana M. Duarte, et al. 2024. “Expansion in Situ Genome Sequencing Links Nuclear Abnormalities to Hotspots of Aberrant Euchromatin Repression.” BioRxiv.

Larson, Adam G., Daniel Elnatan, Madeline M. Keenen, Michael J. Trnka, Jonathan B. Johnston, Alma L. Burlingame, David A. Agard, Sy Redding, and Geeta J. Narlikar. 2017. “Liquid Droplet Formation by HP1α Suggests a Role for Phase Separation in Heterochromatin.” Nature 547: 236–40. 10.1038/nature22822.

Li, J., A. Dong, Kamola Saydaminova, Hill Chang, Guanshi Wang, Hiroshi Ochiai, Takashi Yamamoto, and Alexandros Pertsinidis. 2019. “Single-Molecule Nanoscopy Elucidates RNA Polymerase II Transcription at Single Genes in Live Cells.” Cell 178: 491–506. 10.1016/j.cell.2019.05.029.

Li, Yue, Vasundhara Agrawal, Ranya K.A. Virk, Eric Roth, Wing Shun Li, Adam Eshein, Jane Frederick, et al. 2022. “Analysis of Three-Dimensional Chromatin Packing Domains by Chromatin Scanning Transmission Electron Microscopy (ChromSTEM).” Scientific Reports 12 (1): 1–15. 10.1038/s41598-022-16028-2.

Liu, Zhiyuan, Yujie Chen, Qimin Xia, Menghan Liu, Heming Xu, Yi Chi, Yujing Deng, and Dong Xing. 2023. “Linking Genome Structures to Functions by Simultaneous Single-Cell Hi-C and RNA-Seq.” Science (New York, N.Y.) 380: 1070–76. 10.1126/science.adg3797.

Lombardi, Maria L., Diana E. Jaalouk, Catherine M. Shanahan, Brian Burke, Kyle J. Roux, and Jan Lammerding. 2011. “The Interaction between Nesprins and Sun Proteins at the Nuclear Envelope Is Critical for Force Transmission between the Nucleus and Cytoskeleton.” Journal of Biological Chemistry 286 (30): 26743–53. 10.1074/jbc.M111.233700.

Lorber, Dana, Ron Rotkopf, and Talila Volk. 2020. “A Minimal Constraint Device for Imaging Nuclei in Live: Drosophila Contractile Larval Muscles Reveals Novel Nuclear Mechanical Dynamics.” Lab on a Chip 20 (12): 2100–2112. 10.1039/d0lc00214c.

Lorber, Dana, and Talila Volk. 2022. “Evaluation of Chromatin Mesoscale Organization Evaluation of Chromatin Mesoscale Organization.” APL Bioengineering 010902 (6). 10.1063/5.0069286.

Manzo, Stefano Giustino, Lise Dauban, and Bas van Steensel. 2022. “Lamina-Associated Domains: Tethers and Looseners.” Current Opinion in Cell Biology 74: 80–87. 10.1016/j.ceb.2022.01.004.

Marano, Nicholas, and James M. Holaska. 2025. “The Role of Inner Nuclear Membrane Protein Emerin in Myogenesis.Pdf.” FASEB Journal.

Marcelot, Agathe, Ambre Petitalot, Virginie Ropars, Marie Hélène Le Du, Camille Samson, Stevens Dubois, Guillaume Hoffmann, et al. 2021. “Di-Phosphorylated BAF Shows Altered Structural Dynamics and Binding to DNA, but Interacts with Its Nuclear Envelope Partners.” Nucleic Acids Research 49 (7): 3841–55. 10.1093/nar/gkab184.

Mattout-Drubezki, A., and Y. Gruenbaum. 2003. “Dynamic Interactions of Nuclear Lamina Proteins with Chromatin and Transcriptional Machinery.” Cellular and Molecular Life Sciences 60 (10): 2053–63. 10.1007/s00018-003-3038-3.

Méjat, Alexandre, and Tom Misteli. 2010. “LINC Complexes in Health and Disease LIN.” Nucleus 1 (1): 40–52. 10.4161/nucl.1.1.10530.

Mellad, Jason A, Derek T Warren, and Catherine M Shanahan. 2011. “Nesprins LINC the Nucleus and Cytoskeleton.” Current Opinion in Cell Biology 23: 47–54. 10.1016/j.ceb.2010.11.006.

Miron, Ezequiel, Roel Oldenkamp, Jill M. Brown, David M.S. Pinto, C. Shan Xu, Ana R. Faria, Haitham A. Shaban, et al. 2020. “Chromatin Arranges in Chains of Mesoscale Domains with Nanoscale Functional Topography Independent of Cohesin.” Science Advances 6 (39). 10.1126/sciadv.aba8811.

Misteli, Tom. 2020. “The Self-Organizing Genome: Principles of Genome Architecture and Function.” Cell 183 (1): 28–45. 10.1016/j.cell.2020.09.014.

Nagashima, Ryosuke, Kayo Hibino, S. S. Ashwin, Michael Babokhov, Shin Fujishiro, Ryosuke Imai, Tadasu Nozaki, et al. 2019. “Single Nucleosome Imaging Reveals Loose Genome Chromatin Networks via Active RNA Polymerase II.” Journal of Cell Biology 218 (5): 1511–30. 10.1083/jcb.201811090.

Nozaki, Tadasu, Ryosuke Imai, Mai Tanbo, Ryosuke Nagashima, Sachiko Tamura, Tomomi Tani, Yasumasa Joti, et al. 2017. “Dynamic Organization of Chromatin Domains Revealed by Super-Resolution Live-Cell Imaging.” Molecular Cell 67 (2): 282–293.e7. 10.1016/J.MOLCEL.2017.06.018.

Nozaki, Tadasu, Soya Shinkai, Satoru Ide, Koichi Higashi, Sachiko Tamura, Masa A. Shimazoe, Masaki Nakagawa, et al. 2023. “Condensed but Liquid-like Domain Organization of Active Chromatin Regions in Living Human Cells.” Science Advances 9: eadf1488. 10.1126/sciadv.adf1488.

Ohta, Takao, and Kyozi Kawasaki. 1986. “Equilibrium Morphology of Block Copolymer Melts.” Macromolecules 19 (10): 2621–32.

Oji, Asami, Linda Choubani, Hisashi Miura, and Ichiro Hiratani. 2024. “Structure and Dynamics of Nuclear A / B Compartments and Subcompartments.” Current Opinion in Cell Biology 90: 102406 This.

Padeken, Jan, Stephen P Methot, and Susan M Gasser. 2022. “Establishment of H3K9-Methylated Heterochromatin and Its Functions in Tissue Differentiation and Maintenance.” Nature Reviews Molecular Cell Biology 23: 623. 10.1038/s41580-022-00483-w.

Pancholi, Agnieszka, Tim Klingberg, Weichun Zhang, Roshan Prizak, Irina Mamontova, Amra Noa, Marcel Sobucki, et al. 2021. “RNA Polymerase II Clusters Form in Line with Surface Condensation on Regulatory Chromatin.” Molecular Systems Biology, 1–26.

Pedregosa, Fabian, Gaël Varoquaux, Alexandre Gramfort, Vincent Michel, Bertrand Thirion, Olivier Grisel, Mathieu Blondel, et al. 2011. “Scikit-Learn : Machine Learning in Python.” Jornal of Machine Learning Research 12: 2825–30.

Peng, Ting, Yingping Hou, Haowei Meng, Yong Cao, Xiaotian Wang, Lumeng Jia, Qing Chen, et al. 2023. “Mapping Nucleolus-Associated Chromatin Interactions Using Nucleolus Hi-C Reveals Pattern of Heterochromatin Interactions.” Nature Communications 14: 1–15. 10.1038/s41467-023-36021-1.

Peric-hupkes, Daan, Wouter Meuleman, Ludo Pagie, Sophia W M Bruggeman, Irina Solovei, Wim Brugman, Stefan Graf, et al. 2010. “Molecular Maps of the Reorganization of Genome-Nuclear Lamina Interactions during Differentiation.” Molecular Cell 38: 603–13. 10.1016/j.molcel.2010.03.016.

Piccus, Rachel, and Daniel Brayson. 2020. “The Nuclear Envelope: LINCing Tissue Mechanics to Genome Regulation in Cardiac and Skeletal Muscle.” Biology Letters 16. 10.1098/rsbl.2020.0302.

Plimpton, Steve. 1995. “Fast Parallel Algorithms for Short-Range Molecular Dynamics.” Journal of Computational Physics 117 (1): 1–19. 10.1006/jcph.1995.1039.

Poleshko, Andrey, Cheryl L. Smith, Son C. Nguyen, Priya Sivaramakrishnan, Karen G. Wong, John Isaac Murray, Melike Lakadamyali, Eric F. Joyce, Rajan Jain, and Jonathan A. Epstein. 2019. “H3k9me2 Orchestrates Inheritance of Spatial Positioning of Peripheral Heterochromatin through Mitosis.” ELife 8: 1–24. 10.7554/eLife.49278.

Rajgor, Dipen, and Catherine M. Shanahan. 2013. “Nesprins: From the Nuclear Envelope and Beyond.” Expert Reviews in Molecular Medicine 15 (July): e5. 10.1017/erm.2013.6.

Razafsky, David, Denis Wirtz, and Didier Hodzic. 2014. “Nuclear Envelope in Nuclear Positioning and Cell Migration.” Advances in Experimental Medicine and Biology, 471–90. 10.1007/978-1-4899-8032-8_21.

Ricci, Maria Aurelia, Carlo Manzo, María Filomena García-Parajo, Melike Lakadamyali, and Maria Pia Cosma. 2015. “Chromatin Fibers Are Formed by Heterogeneous Groups of Nucleosomes in Vivo.” Cell 160 (6): 1145–58. 10.1016/j.cell.2015.01.054.

Rippe, Karsten, and Argyris Papantonis. 2022. “Functional Organization of RNA Polymerase II in Nuclear Subcompartments.” Current Opinion in Cell Biology 74: 88–96. 10.1016/j.ceb.2022.01.005.

Robson, Michael I., Alessa R. Ringel, and Stefan Mundlos. 2019. “Regulatory Landscaping: How Enhancer-Promoter Communication Is Sculpted in 3D.” Molecular Cell. 10.1016/j.molcel.2019.05.032.

Rothballer, Andrea, and Ulrike Kutay. 2013. “The Diverse Functional LINCs of the Nuclear Envelope to the Cytoskeleton and Chromatin.” Chromosoma 122 (5): 415–29. 10.1007/s00412-013-0417-x.

Salari, Hossein, Geneviève Fourel, and Daniel Jost. 2024. “Transcription Regulates the Spatio-Temporal Dynamics of Genes through Micro-Compartmentalization.” Nature Communications 15. https://www.biorxiv.org/content/10.1101/2023.07.18.549489v2 %0A https://www.biorxiv.org/content/10.1101/2023.07.18.549489v2.abstract.

Sanders, Jacob T., Rosela Golloshi, Priyojit Das, Yang Xu, Peyton H. Terry, Darrian G. Nash, Job Dekker, and Rachel Patton McCord. 2022. “Loops, Topologically Associating Domains, Compartments, and Territories Are Elastic and Robust to Dramatic Nuclear Volume Swelling.” Scientific Reports 12: 1–19. 10.1038/s41598-022-08602-5.

Sanulli, S, M J Trnka, V. Dharmarajan, R. W. Tibble, B. D. Pascal, A. L. Burlingame, P. R. Griffin, J. D. Gross, and G. J. Narlikar. 2019. “HP1 Reshapes Nucleosome Core to Promote Phase Separation of Heterochromatin.” Nature 575 (November). 10.1038/s41586-019-1669-2.

Sazer, Shelley, and Helmut Schiessel. 2018. “The Biology and Polymer Physics Underlying Large-Scale Chromosome Organization.” Traffic 19 (2): 87–104. 10.1111/tra.12539.

Sears, Rhiannon M, and Kyle J Roux. 2020. “Diverse Cellular Functions of Barrier-to-Autointegration Factor and Its Roles in Disease.” Journal of Cell Science 133. 10.1242/jcs.246546.

See, Kelvin, Yemin Lan, Joshua Rhoades, Rajan Jain, Cheryl L. Smith, and Jonathan A. Epstein. 2019. “Lineage-Specific Reorganization of Nuclear Peripheral Heterochromatin and H3K9Me2 Domains.” Development (Cambridge) 146 (3). 10.1242/dev.174078.

Shah, Parisha P., Garrett T. Santini, Kaitlyn M. Shen, and Rajan Jain. 2023. “InterLINCing Chromatin Organization and Mechanobiology in Laminopathies.” Current Cardiology Reports 25: 307–14. 10.1007/s11886-023-01853-2.

Shao, Wen, Xianju Bi, Yixuan Pan, Boyang Gao, Jun Wu, Yafei Yin, Zhimin Liu, et al. 2022. “Phase Separation of RNA-Binding Protein Promotes Polymerase Binding and Transcription.” Nature Chemical Biology 18: 70–80. 10.1038/s41589-021-00904-5.

Silva, Shanelle De, Zhijuan Fan, Baoqiang Kang, Catherine M Shanahan, and Qiuping Zhang. 2023. “Nesprin-1 : Novel Regulator of Striated Muscle Nuclear Positioning and Mechanotransduction.” Biochemical Society Transactions 51 (3): 1331–45.

Sobo, Joan M, Nicholas S Alagna, Sean X Sun, Katherine L Wilson, and Karen L Reddy. 2024. “Lamins : The Backbone of the Nucleocytoskeleton Interface.” Current Opinion in Cell Biology 86: 1–16.

Sood, Varun, and Tom Misteli. 2022. “The Stochastic Nature of Genome Organization and Function.” Current Opinion in Genetics and Development 72: 1–8. 10.1016/j.gde.2021.10.004.

Sosa, Brian A., Andrea Rothballer, Ulrike Kutay, and Thomas U. Schwartz. 2012. “LINC Complexes Form by Binding of Three KASH Peptides to Domain Interfaces of Trimeric SUN Proteins.” Cell 149: 1035–47. 10.1016/j.cell.2012.03.046.

Steensel, Bas van, and Andrew S. Belmont. 2017. “Lamina-Associated Domains: Links with Chromosome Architecture, Heterochromatin, and Gene Repression.” Cell 169 (5): 780–91. 10.1016/j.cell.2017.04.022.

Strom, Amy R., Alexander V. Emelyanov, Mustafa Mir, Dmitry V. Fyodorov, Xavier Darzacq, and Gary H. Karpen. 2017. “Phase Separation Drives Heterochromatin Domain Formation.” Nature 547: 241–45. 10.1038/nature22989.

Szabo, Quentin, Daniel Jost, Jia Ming Chang, Diego I. Cattoni, Giorgio L. Papadopoulos, Boyan Bonev, Tom Sexton, et al. 2018. “TADs Are 3D Structural Units of Higher-Order Chromosome Organization in Drosophila.” Science Advances 4: 1–13. 10.1126/sciadv.aar8082.

Tapley, Erin C., and Daniel A. Starr. 2013. “Connecting the Nucleus to the Cytoskeleton by SUN-KASH Bridges across the Nuclear Envelope.” Current Opinion in Cell Biology 25: 57–62. 10.1016/j.ceb.2012.10.014.

The pandas development team. 2020. “Pandas-Dev/Pandas: Pandas.” Zenodo. February 2020. 10.5281/zenodo.18675244.

Thompson, Aidan P, H Metin Aktulga, Richard Berger, Dan S Bolintineanu, W Michael Brown, Paul S Crozier, Pieter J. in ’t Veld, et al. 2022. “LAMMPS - a Flexible Simulation Tool for Particle-Based Materials Modeling at the Atomic, Meso, and Continuum Scales.” Computer Physics Communications 271 271: 108171.

Towbin, Benjamin D, Cristina Gonzalez-Aguilera, Ragna Sack, Dimos Gaidatzis, Veronique Kalck, Peter Meister, Peter Askjaer, and Susan M Gasser. 2012. “Step-Wise Methylation of Histone H3K9 Positions Heterochromatin at the Nuclear Periphery.” Cell 150: 934–47.

Trojanowskia, Jorge, and Karsten Rippe. 2022. “Transcription Factor Binding and Activity on Chromatin.” Current Opinion in Structural Biology 31.

Tullius, Thomas W., R. Stefan Isaac, Danilo Dubocanin, Jane Ranchalis, L. Stirling Churchman, and Andrew B. Stergachis. 2024. “RNA Polymerases Reshape Chromatin Architecture and Couple Transcription on Individual Fibers.” Molecular Cell 84: 3209–22. 10.1016/j.molcel.2024.08.013.

Ulianov, Sergey V., Semen A. Doronin, Ekaterina E. Khrameeva, Pavel I. Kos, Artem V. Luzhin, Sergei S. Starikov, Aleksandra A. Galitsyna, et al. 2019. “Nuclear Lamina Integrity Is Required for Proper Spatial Organization of Chromatin in Drosophila.” Nature Communications 10: 1–11. 10.1038/s41467-019-09185-y.

Uzer, Gunes, Clinton T Rubin, and Janet Rubin. 2016. “Cell Mechanosensitivity Is Enabled by the LINC Nuclear Complex.” Current Molecular Biology Reports 2 (1): 36–47. 10.1007/s40610-016-0032-8.Cell.

Verschure, Pernette J., Ineke van der Kraan, Erik M.M. Manders, Deborah Hoogstraten, Adriaan B. Houtsmuller, and Roel van Driel. 2003. “Condensed Chromatin Domains in the Mammalian Nucleus Are Accessible to Large Macromolecules.” EMBO Reports 4 (9): 861–66. 10.1038/sj.embor.embor922.

Virtanen, Pauli, Ralf Gommers, Travis E. Oliphant, Matt Haberland, Tyler Reddy, David Cournapeau, Evgeni Burovski, et al. 2020. “SciPy 1.0: Fundamental Algorithms for Scientific Computing in Python.” Nature Methods 17 (March): 261–72. 10.1038/s41592-019-0686-2.

Wagh, Kaustubh, Momoko Ishikawa, David A. Garcia, Diana A. Stavreva, Arpita Upadhyaya, and Gordon L. Hager. 2021. “Mechanical Regulation of Transcription: Recent Advances.” Trends in Cell Biology 31 (6): 457–72. 10.1016/j.tcb.2021.02.008.

Walt, Stefan van der, Johannes L. Schönberger, Juan Nunez-Iglesias, François Boulogne, Joshua D. Warner, Neil Yager, Emmanuelle Gouillart, Tony Yu, and Scikit-image Contributors. 2014. “Scikit-Image: Image Processing in Python.” PeerJ, 1–18. 10.7717/peerj.453.

Wang, Baihui, Qiang Luo, and Ohad Medalia. 2025. “Lamins and Chromatin Join Forces.” Advances in Biological Regulation 95.

Wang, Shuoshuo, Elizabeth Stoops, C. P. Unnikannan, Barak Markus, Adriana Reuveny, Elly Ordan, and Talila Volk. 2018. “Mechanotransduction via the LINC Complex Regulates DNA Replication in Myonuclei.” Journal of Cell Biology 217 (6): 2005–18. 10.1083/jcb.201708137.

Wei, Ming Tzo, Yi Che Chang, Shunsuke F. Shimobayashi, Yongdae Shin, Amy R. Strom, and Clifford P. Brangwynne. 2020. “Nucleated Transcriptional Condensates Amplify Gene Expression.” Nature Cell Biology 22 (10): 1187–96. 10.1038/s41556-020-00578-6.

Wong, Xianrong, Tsui-han Loo, and Colin L Stewart. 2021. “LINC Complex Regulation of Genome Organization and Function.” Current Opinion in Genetics & Development 67: 130–41.

Yerima, Ghafar, Nya Domkam, Jessica Ornowski, Zeinab Jahed, and Mohammad R.K. Mofrad. 2023. “Force Transmission and SUN-KASH Higher-Order Assembly in the LINC Complex Models.” Biophysical Journal 122 (23): 4582–97. 10.1016/j.bpj.2023.11.001.

Zenk, Fides, Yinxiu Zhan, Pavel Kos, Eva Löser, Nazerke Atinbayeva, Melanie Schächtle, Guido Tiana, Luca Giorgetti, and Nicola Iovino. 2021. “HP1 Drives de Novo 3D Genome Reorganization in Early Drosophila Embryos.” Nature 593 (May).

Zhang, Shu, Nadine Übelmesser, Mariano Barbieri, and Argyris Papantonis. 2023. “Enhancer – Promoter Contact Formation Requires RNAPII and Antagonizes Loop Extrusion.” Nature Genetics 55 (May). 10.1038/s41588-023-01364-4.

Zhang, Shu, Nadine Übelmesser, Natasa Josipovic, Giada Forte, Johan A. Slotman, Michael Chiang, Henrike Johanna Gothe, et al. 2021. “RNA Polymerase II Is Required for Spatial Chromatin Reorganization Following Exit from Mitosis.” Science Advances 7 (43): 1–14. 10.1126/sciadv.abg8205.

Zhou, Can, Li Rao, Catherine M Shanahan, and Qiuping Zhang. 2018. “Nesprin-1/2:Roles in Nuclear Envelope Organisation, Myogenesis and Muscle Disease.” Biochemical Society Transactions 46: 311–20.

